# Localization of *Drosophila* formin, Cappuccino, influences posterior oocyte organization

**DOI:** 10.1101/2024.07.22.604638

**Authors:** Hannah M. Bailey, Peter B. M. Cullimore, Liam A. Bailey, Margot E. Quinlan

## Abstract

Cappuccino (Capu) and Spire build actin networks in numerous systems, including the mouse oocyte, melanocytes, and the Drosophila oocyte. As observed in mammalian systems, the localization of the Capu homologues (FMN1/2), influences the function of the actin network. Therefore, we established and interrogated the impact of altering Capu’s localization in the Drosophila oocyte to better understand its role and that of the actin mesh it builds. This mesh restricts bulk cytoplasmic flows, streaming, but otherwise remains undescribed functionally. Using a gene specific driver, capu-Gal4, to better study Capu transgenes, we found that fertility was markedly decreased when restricting Capu to membranes in the oocyte, although its canonical role in actin mesh assembly was apparently unaltered. Instead, we observed a defect in posterior anchoring of the mRNA oskar during mid-oogenesis. However, the defect did not fall into the traditional posterior group phenotype. The data suggest that Capu, independently of Spire, tethers the posterior determinants to the cortex but does not anchor them to each other, supporting that Capu localization influences the posterior oocyte organization.

## Introduction

*Drosophila* oogenesis has long served as a model for studying polarization of developing cells. Careful regulation of the actin cytoskeleton and a range of actin-based structures are essential to cell polarity establishment and maintenance. Among these structures, the actin mesh, is poorly understood. It is a network of filamentous actin, that fills the developing oocyte specifically during developmental stages 5-10A (Dahlgaard et al., 2007). If the network is not built or persists beyond stage 10B, fertility is severely decreased (Bor et al., 2015; Dahlgaard et al., 2007; Quinlan, 2013). In both cases, polarized localization of mRNAs is disrupted. It is thought that the actin mesh indirectly impacts mRNA localization by restricting microtubule-dependent cytoplasmic streaming (Dahlgaard et al., 2007; Gutzeit and Koppa, 1982). Two proteins, specifically actin nucleators, Spire (Spir) (Quinlan et al., 2005) and Cappuccino (Capu) (Emmons et al., 1995) are required for proper formation of this meshwork (Dahlgaard et al., 2007). Their direct interaction results in synergistic actin assembly, which is required to build the mesh (Bradley et al., 2019; Quinlan, 2013). It follows that mutation of either gene leads to absence/loss of mesh (Dahlgaard et al., 2007; Manseau and Schupbach, 1989). Downstream of mesh loss, premature onset of fast cytoplasmic streaming, loss of cell polarity, and infertility are observed. It is difficult to determine if the sole consequence of *spir* or *capu* mutation is loss of the actin mesh or if other processes are disrupted. Premature streaming is so severe that it may be masking other processes that require the mesh, Spir, and/or Capu. In fact, there exists evidence of other roles for Spir and Capu have been presented (Alzahofi et al., 2020; Chang et al., 2011; Scheffler et al., 2021; Schuh and Ellenberg, 2008; Stürner et al., 2022). Here we present evidence that Capu plays at least one alternate role in the Drosophila oocyte, which may be independent of Spir.

In the developing mouse oocyte, an analogous actin meshwork is built by mammalian homologs of Spir (Spire-1 and Spire-2) and Capu (FMN-2) (Pfender et al., 2011; Schuh and Ellenberg, 2008). FMN2 and Spire-1/2 colocalize on Rab11-positive vesicles and the cortex, where they nucleate actin filaments. Myosin V (MyoV) movement along this network of actin generates a pushing force – contraction – to centrally position the mitotic spindle and transport Rab11-positive vesicles in meiosis (Almonacid and Verlhac, 2021; Holubcová et al., 2013; Schuh and Ellenberg, 2008). Interestingly, the mammalian paralog, FMN1, is cytosolic in mouse melanocytes, which have Spire-1/2 abutting Rab27-positive melanosomes. In this case, MyoV drives dispersion of melanosomes, as opposed to contraction (Alzahofi et al., 2020).

The implication of the mouse oocyte and melanocyte models is that localization of the formin defines the structure built by Spir and Capu, thereby determining the consequences of their activity. Spir is enriched at the cortex in the *Drosophila* oocyte (Quinlan 2007, 2013). All Spir proteins contain a mFYVE domain, which binds negatively charged membranes, consistent with its presence on vesicles and in the cortex (Kerkhoff et al., 2001; Tittel et al., 2015). Based on transgene expression, Capu appears to be diffuse in the developing *Drosophila* oocyte, though it is enriched at the cortex of nurse cells (Bor et al., 2015; Dahlgaard et al., 2007; Quinlan, 2013). This localization initially surprised us based on the mouse oocyte data and the fact that Capu directly interacts with membrane-associated Spir to function (Quinlan, 2013). Therefore, we sought to determine the localization of endogenous protein and study the effect of altering Capu localization in the *Drosophila* oocyte.

A notable difference between FMN1 and FMN2 is the presence of an N-terminal glycine only in FMN2 that is predicted to be N-myristoylated by n-myristoyltransferase (NMT). Myristoylation may drive FMN2 to membranes, as observed in the mouse oocyte. Nine unique isoforms of Cappuccino are predicted in the fly (FlyAtlas2, FlyBase)(Leader et al., 2018; Öztürk-Çolak et al., 2024). A closer look at these isoforms reveals a myristoylation site in the N-terminal exon of isoforms D/E/F/J. CapuA is the canonical isoform expressed in the ovary. It does not have a predicted myristoylation site. The diffuse localization of CapuA, suggests that Spir, Capu, and the mesh may function more like the melanocyte – important for dispersion - than the mouse oocyte – driving contractile forces.

We created endogenously tagged genes to establish high resolution localization data for Spir and Capu. We then tested the impact of altering Capu localization, by utilizing transgene rescue with a new gene-specific driver. These experiments enabled us to identify an alternate role for Capu in oogenesis, linking a structure critical for organization of posterior determinants to the oocyte cortex, which appears to be independent of Spir.

## Results

### Capu localization

Because function is necessarily linked to localization, we decided to revisit the question of where Capu is found in the Drosophila oocyte. Previous work relied on transgene expression because multiple attempts by multiple groups to generate a specific Capu antibody for immunofluorescence were unsuccessful. The *capu* gene has five potential start sites and 9 transcripts, but all of the splice variants share their C-termini. So, we took advantage of a MiMIC line (MI05735) to insert a C-terminal tag in the endogenous gene (Fig. S1A). As proof of principle, we first inserted a duplicate of the remaining gene, such that a full-length untagged version of Capu would be expressed under the same control as wild type. The *capu* insertion worked well, resulting in 83% fertility (Fig. S1B). We then inserted tags using the same method. We started with GFP because transgene rescue with Capu-GFP has been successful (Quinlan 2013). Homozygous females expressing Capu-GFP were fertile at levels comparable to those with the untagged gene, 80% (Fig. S1B). Our efforts to image Capu-GFP in live samples or by immunofluorescence suggested that Capu is expressed at low levels (Fig. S1B,C, a-a”). Immunofluorescence revealed that Capu is largely diffuse in the oocyte, as observed with transgenes. We, therefore, inserted mScarlet-3XOLLAS (Fig. S1B, C, b-b”). The localization patterns detected with GFP and OLLAS are consistent with one another and similar to previously reported transgene expression (Dahlgaard et al., 2007; Quinlan, 2013; Rosales-Nieves et al., 2006). Broadly, Capu is diffuse throughout the egg chamber and enriched in the nurse cell cortex. Interestingly, we observed Capu enriched at the oocyte cortex during mid oogenesis, when the mesh is present (stages 6-9) (Fig. S1C). However, we only observed cortical enrichment in the oocyte in fixed samples. In addition, we observed Capu in specialized follicle cells: the migrating border cell cluster (Fig. S1C, a’ arrow) and posterior polar cells (Fig. S1C, b” arrow). Experiments using germline-specific drivers could not have revealed these previously undescribed localizations of Capu.

### A *capu*-GAL4 driver

In the past, Capu transgene expression was usually driven by *nanos-*Gal4-vp16 (*nos*-Gal4) and always by germline specific drivers (Dahlgaard et al., 2007; Quinlan, 2013; Rosales-Nieves et al., 2006). While rescue of fertility in the null is observed, it is incomplete. For example, when driving CapuA or GFP-CapuA, fertility was 56 and 36%, respectively (Quinlan, 2013). Low fertility could reflect a requirement for alternate splice variants of Capu or a mismatch in Capu’s spatiotemporal expression pattern when driven by *nos*-Gal4. For example, *nos*-Gal4 does not drive expression in the border and polar cells. To test these possibilities, we built a gene-specific driver, *capu-*Gal4-K10 (*capu-* Gal4). We inserted Gal4-K10 at the MI05737 landing site within *capu,* using a modified Trojan-Gal4 design strategy (Lee et al., 2018; Venken et al., 2011). Expression driven by *capu-*Gal4 begins as early as stage 2A in the germarium, increases in intensity at stage one and is continually expressed until the onset of late oogenesis (Fig. 1C), a pattern that markedly differs from *nos*-Gal4 (Hudson and Cooley, 2014). Using *capu-*Gal4 driven CapuA-GFP to rescue a *capu* null background resulted in 90% fertility: a great improvement over *nos*-Gal4 driven rescue experiments (Table 1). Localization of CapuA matches that seen when the gene is driven by *nos*-Gal4: largely diffuse with enrichment at the nurse cell cortex (Fig. S1). We do not see follicle cell expression, presumably due to the 3’-UTRs used in the Gal4 driver and the pUASp insertion. Thus, we conclude that primarily germline expression of CapuA is sufficient to rescue fertility, consistent with expression data indicating that it is the major isoform expressed in the ovary (Leader et al., 2018). Further, the process is sensitive to the timing of Capu expression. We, therefore, use only *capu*-Gal4 as a driver in the following studies.

**Figure 1.**
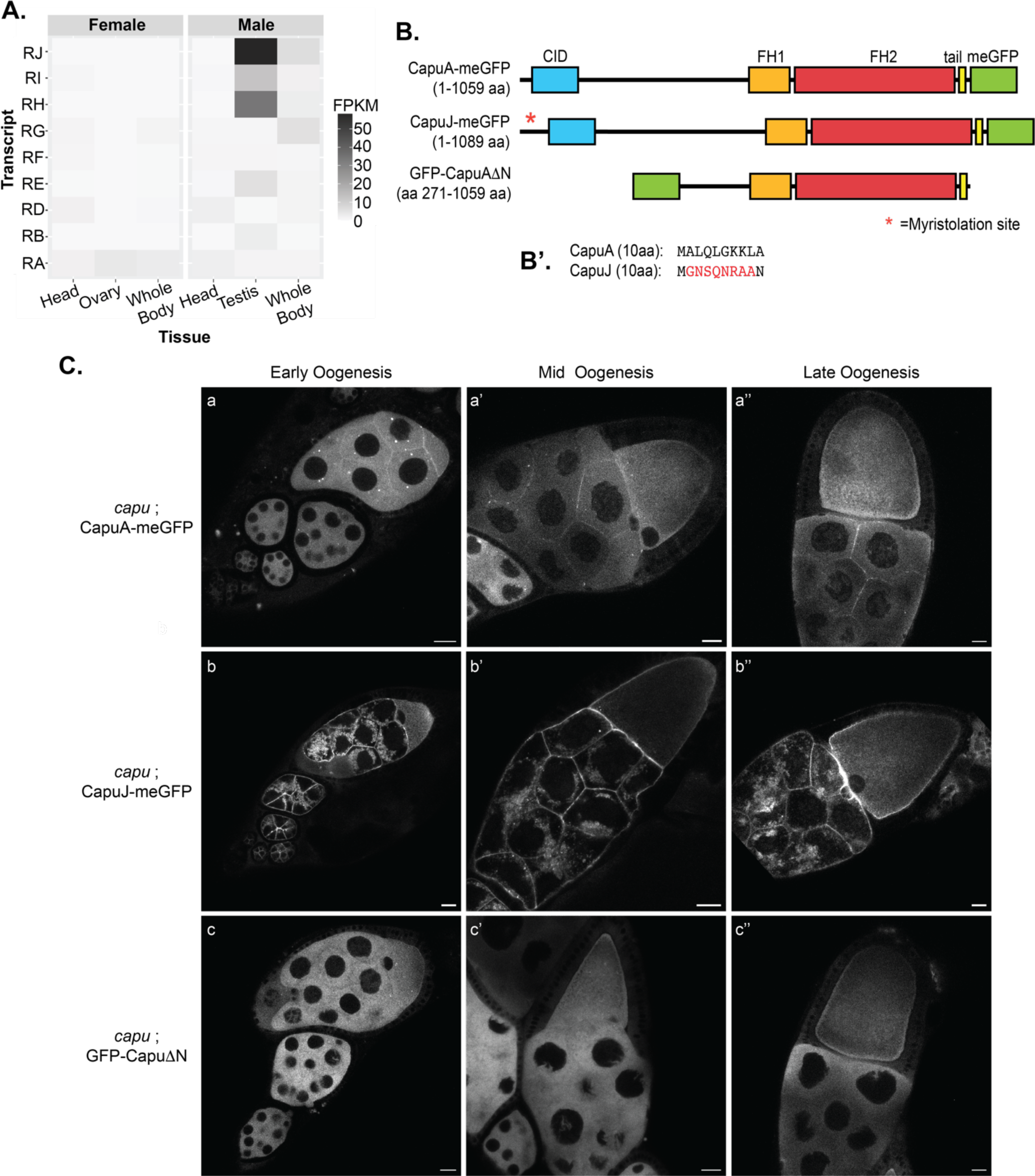
Myristoylation of Capu alters localization. (A) Heatmap of transcript expression data from FlyAtlas2, modified from Leader et al, 2018. (B) Domain maps of transgenes used in this study. CapuA-GFP, CapuJ-GFP, and GFP-CapuAΔN. Numbers in the parentheses are amino acids of the constructs, not including the GFP. Major domains indicated are; CID, Capu inhibitory domain (blue); FH1, formin homology 1 (orange); FH2, formin homology 2 (red); tail (yellow). A green box is included to indicate the location of the GFP. The red asterisk indicates the region of CapuJ where the myristoylation site is found. (B’) The first ten amino acids of CapuA and CapuJ. The predicted myristoylation motif is indicated in red. (C) Capu localization in live egg chambers throughout oogenesis. (a-a”) CapuA localization is enriched at the nurse cell cortex, interface between the nurse cell and oocyte, and is largely diffuse in nurse cell and oocyte cytoplasm. (b-b”) CapuJ exhibits an increased enrichment at the nurse cell cortexes, and nurse cell/oocyte interface. Cytoplasmic diffuse localization is lost in the nurse cells and there is a stronger cortical localization pattern in the oocyte during mid and late oogenesis. (c-c”) CapuΔN localization is completely diffuse through the nurse cells and oocyte for all developmental stages. Scale bars: 20 μm.

**Table 1:** Table of fertility data from genetic rescue experiments. The genetic background is in parentheses, all transgenes are under UASp regulation. The % hatched is reported as the average of three independent trials, with the number counted being the sum total eggs collected from all trials. Fertility is reported as the percentage of eggs that hatched within 24 hours of being laid. All test female flies were crossed to wild-type males to evaluate the maternal contribution to the rescue rate.

| Transgene<br>(Df(2L)ed<sup>1</sup>/capu-Gal4-K10) | % Hatched | Number counted |
| --- | --- | --- |
| UASp::CapuA-meGFP | 89.5 | 637 |
| UASp::CapuJ-meGFP | 47.8 | 694 |
| UASp::eGFP-CapuΔN | 43.1 | 800 |
| Null/No Rescue | 0.66 | 598 |

### Myristoylation of Capu alters localization and decreases fertility

Capu isoforms D/E/F/J are highly expressed in the male testis but not detected in the female ovary (Fig. 1A)(Leader et al., 2018). As noted above, these isoforms share their first coding exon containing a predicted N-terminal myristoylation site, which could target Capu to membranes. We reasoned that by expressing one of these isoforms we could alter Capu localization in the oocyte. We selected CapuJ because the coding regions of CapuA and CapuJ only differ by their most N-terminal exons (Benson et al., 2012) (Fig. 1B-B’).

CapuJ-GFP is largely membrane bound (Fig. 1C). Higher magnification imaging reveals strong cortical localization in the oocyte as well as punctate-like signal in the cytoplasm (Fig. S1D). In addition, there is an aggregate-like localization pattern in the nurse cell cytoplasm and an increase in signal at the nurse cell-oocyte interface (Fig. 1C, b-b”). Based on the altered localization, we conclude that CapuJ is myristoylated when expressed in the *Drosophila* egg chamber.

We also considered GFP-CapuΔN, a truncated version of CapuA, which lacks the N-terminus through its autoinhibitory domain (CID; Fig. 1B). GFP-CapuΔN was previously shown to rescue mesh formation and *oskar* localization when driven by *nos-*Gal4 (Dahlgaard et al., 2007). Live imaging of GFP-CapuΔN driven by *capu*-Gal4 reveals localization patterns similar to those previously reported with *nos*-Gal4 (Dahlgaard et al., 2007; Quinlan, 2013). GFP-CapuΔN is completely diffuse; no cortical enrichment is detected in the nurse cells or at the nurse cell-oocyte interface (1C, c-c”).

As the spatial regulation is critical for formin function in other systems (Alzahofi et al., 2020; Azoury et al., 2008; Holubcová et al., 2013), we asked whether altering the localization in the *Drosophila* egg chamber influences Capu function and, therefore, fertility. Interestingly, CapuJ-meGFP-expressing flies were only 48% fertile. In addition, GFP-CapuΔN expressing flies were 43% fertile (Table 1). These marked decreases in fertilty demonstrate that Capu’s function is impaired when the N-terminus is altered, perhaps due to changes in localization.

### CapuJ builds an actin mesh

A straightforward explanation for the decreased fertility with CapuJ rescue would be that the actin mesh is misregulated. To be assembled, the actin mesh requires direct interaction between Spir and Capu (Bradley et al., 2019; Dahlgaard et al., 2007; Quinlan, 2013). By restricting Capu’s localization within the oocyte, Capu’s actin assembly activity and synergy with Spir could be disrupted. We stained for the actin mesh in rescue egg chambers and found no difference in mesh presence (mid-oogenesis) or the timing of removal (late oogenesis) (Fig. 2A). As previously reported (Dahlgaard et al., 2007) GFP-CapuΔN rescue exhibited formation of a denser actin mesh in the nurse cells. We attribute the apparently increased activity to loss of autoinhibition. Similarly, *nos*-Gal4 driven expression of GFP-CapuΔN, but not GFP-CapuA, markedly decreased fertility when expressed in wild type background (Bor et al., 2015). Thus, both excess actin in nurse cells and hyperactive Capu in the oocyte may be contributors to the decreased fertility in GFP-CapuΔN rescue flies (Table 1). Therefore, we include CapuΔN in some of our studies but interpret it with care when comparing it to CapuA and CapuJ. Altogether these data demonstrate that membrane-bound Capu (CapuJ) retains actin assembly activity and is capable of forming an actin meshwork with wildtype timing, indistinguishable from cytosolic Capu (CapuA).

**Figure 2.**
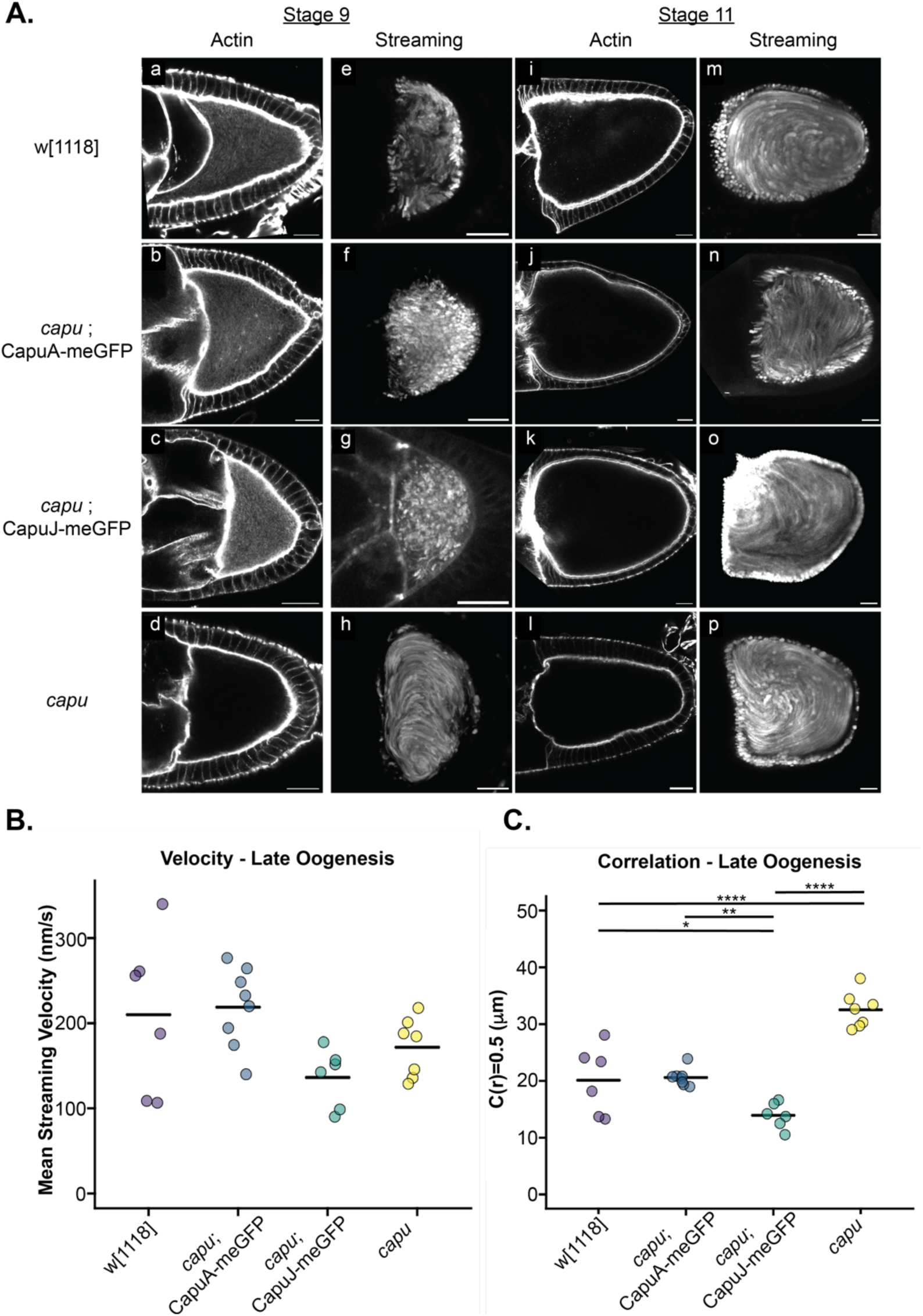
Membrane-bound Capu is sufficient to rescue actin mesh formation and timing. (A.) Stage 9 and 11 egg chambers stained with AlexaFluor488-phalloidin to examine the actin mesh in transgenic flies. For both CapuA (Df(2L)ed^1^/*capu-*Gal4-K10; UASp::CapuA-meGFP/+) and CapuJ (Df(2L)ed^1^/*capu-*Gal4-K10; UASp::CapuJ-meGFP/+) rescue, the actin mesh is properly formed (stage 9, a-d) and removed (stage 11, i-l) during oogenesis with a wildtype (w[1118]) control for comparison. *capu* null (Df(2L)ed^1^/*capu-*Gal4-K10; +/+) included to show failed rescue. Maximum intensity projections of autofluorescent yolk granule motion over a period of 5 minutes for transgenic egg chambers mid-oogenesis (typical stage 9, e-h) and late oogenesis (stage 11, m-p) are shown for each background. Scale bars: 20μm. (B-C) Analysis of streaming velocities and pattern in late oogenesis. (Mid oogenesis shown in Figure S2.) Dot plots of streaming velocities and pattern (correlation radius = 0.5), the bar in each dataset indicates the average value. N ≧ 6 for each stage 11 egg chamber analyzed. (B) Late streaming velocity is not significantly different for any of the genotypes measured (∼200 nm/s), see methods for analysis. (C) Analysis of the late oogenesis streaming pattern, correlation (described in the methods), using a custom code (2DCorrelation, LiamABailey, GitHub). The correlation pattern of streaming during late oogenesis in *capu* null oocytes is significantly increased (∼32 μm, ****, p = 9.87E-06) in comparison to wildtype (20 µm). The CapuJ rescue has a significantly lower correlation pattern (15 μm) in comparison to wildtype (20 μm, *, p = 0.0265), CapuA rescue (21 μm, **, p = 0.00919), and *capu* null (32 μm, ****, p = 1.09E-08). All p values were determined via one-way ANOVA, post-hoc Tukey-Kramer Multiple Comparisons Test. See methods for more details.

To functionally assess the actin mesh, we analyzed the cytoplasmic fluid flows, so-called cytoplasmic streaming, during mid and late oogenesis (Fig. 2A, streaming-h, m-p). We analyzed timelapse images of yolk granule movement to determine velocity (PIVlab, MATLAB) and pattern (2DCorrelation, LiamABailey GitHub). To describe the pattern of motions, we measured the degree of correlation of motion and report the maximal radius at which the streaming topology remains half correlated (C(r)=0.5). No significant difference was observed during mid oogenesis for velocity or correlation between wildtype and rescues, leading us to conclude that the mesh is functional and streaming is properly regulated at this stage (Fig. S2). (Unless otherwise stated, we used one-way ANOVA and post-hoc Tukey-Kramer Multiple Comparisons Tests (TKMC) to determine significance, p=0.05)

We found differences in streaming during late oogenesis. While there was no significant difference in velocity (Fig. 2B), we did find an intriguing difference in the pattern/correlation of fast streaming during late oogenesis (Fig. 2C). In wildtype oocytes, at the (C(r)=0.5) of fast streaming, the radius is ∼20 µm on average. CapuJ rescue oocytes were found to have a significantly lower radius of ∼15 µm. The change is the opposite direction from the correlation pattern of *capu* null oocytes, which have significantly higher correlated radii (C(r)=0.5 radius of ∼32 µm) compared to wildtype (p=1.09E-08). While interesting, the change in pattern correlation is small and we do not predict that it is substantial enough to explain the fertility defect observed with CapuJ rescue (Table 1).

During oogenesis, the known role of Capu is its involvement in actin mesh formation, which regulates cytoplasmic streaming. By rescuing with membrane bound Capu, CapuJ, we have determined that actin mesh assembly and streaming are largely unaltered, leaving the reduction in fertility unexplained. This poses important questions, such as: are there additional roles for Capu during *Drosophila* oogenesis? Further, could a role for the mesh, other than streaming regulation, be disrupted because the mesh organization is altered when CapuJ builds it?

### Restricting Capu to membranes leads to disrupted posterior pole during mid-oogenesis

The actin mesh, microtubule organization and *oskar* localization are tightly correlated (Babu et al., 2003; Dahlgaard et al., 2007; Glotzer et al., 1997; Krauss et al., 2009; Serbus, 2005). In both *spir* and *capu* nulls, localization of polarity factors, including *oskar, nanos*, Vasa, and Staufen, is impaired (Dahlgaard et al., 2007; Ephrussi et al., 1991; Kim-Ha et al., 1991; Manseau and Schupbach, 1989; St. Johnston et al., 1991). A straightforward explanation is that delivery and maintenance fail due to disorganized microtubules and premature fast streaming. If this is the whole story, our analysis of the actin mesh and streaming in CapuJ-GFP-expressing flies would lead to the prediction that localization of polarity factors would not be altered.

The altered localization pattern of CapuJ-GFP, including enrichment at the posterior oocyte cortex (Fig. S1D), and oocyte-nurse cell interface during mid-oogenesis, motivated us to examine both anterior and posterior factors. We used smiFISH (Calvo et al., 2021; Lu et al., 2023; Tsanov et al., 2016) to label polarity determinants, *gurken (grk)* and *bicoid (bcd)* at the anterior oocyte and *oskar (osk)* at the posterior, specifically in mid-to late-stage 9 egg chambers. We observed no difference in mRNA localization at the anterior when compared to wildtype (Fig. S3). This is consistent with previous studies that implicate microtubules in localization and anchoring of *grk* and *bcd* (Jaramillo et al., 2008; Trovisco et al., 2016; Weil et al., 2006).

In contrast, *osk* mRNA at the posterior oocyte is altered in the CapuJ-GFP rescue background (Fig. 3A). In stage 9 wildtype egg chambers *osk* is tightly localized to the posterior cap. Posterior localization of *osk* is completely lost if fast streaming is premature – as observed in *capu* null oocytes (Fig. 3A). When CapuJ-GFP was expressed, we observed a localization pattern that differed from wildtype but was not nearly as dramatic as seen in the *capu* null. To describe and compare the localization patterns we measured the intensity profile of the smiFISH signal around the oocyte cortex (Fig. S4) and along the anterior/posterior axis in the center of the oocyte (Fig. 3B). For all rescue backgrounds the localization of *osk* around the cortex exhibited a greater spread at the posterior (Fig. S4). The most striking difference was observed in the anterior to posterior (AP) localization of *osk*. To characterize these differences, we measured the distance of the intensity peak from the posterior oocyte (distance), the full width of the intensity signal at half max (FWHM) and the amplitude of the intensity signal for each genotype (average amplitude). Wildtype *osk* was closely localized to the posterior oocyte (distance = 0.035, FWHM = 0.054) at high intensity (average amplitude 0.811) (Fig. 3B-B’). CapuA-GFP and GFP-CapuΔN rescue had similar localization patterns to each other, with an increased spread (FWHM was twice that of wildtype, 0.12, Fig. 3B-B’). The peak intensity was also shifted twice as far from the posterior (distance = ∼0.8) with an overall 20% reduction in peak amplitude (average amplitude = ∼0.65) (Fig. 3B-B’). CapuJ-GFP rescue oocytes had the most dramatic changes in the AP localization pattern of *osk* (Fig. 3B-B’). *osk* on average was reduced by 40% in intensity (average amplitude = 0.46) with a fourfold increase in distance from the posterior (distance = 0.15) and threefold increase in spread (FWHM = 0.17) in comparison to wildtype oocytes (Fig. 3B-B’). To determine what was causing the spread in the average peak intensity, we analyzed individual oocyte trace data (Fig. 3C-C’). From this we discerned that the average reduction was due to a significant difference in peak distance from the posterior (Fig. 3C’, p=2.02E-08) as opposed to increased spread of *osk* in individual oocytes (Fig. 3C). This suggests that *osk* mRNA is organized but not tightly anchored at the posterior. It is also consistent with the distance from the posterior, as opposed to cortical spread, contributing more to fertility defects.

**Figure 3.**
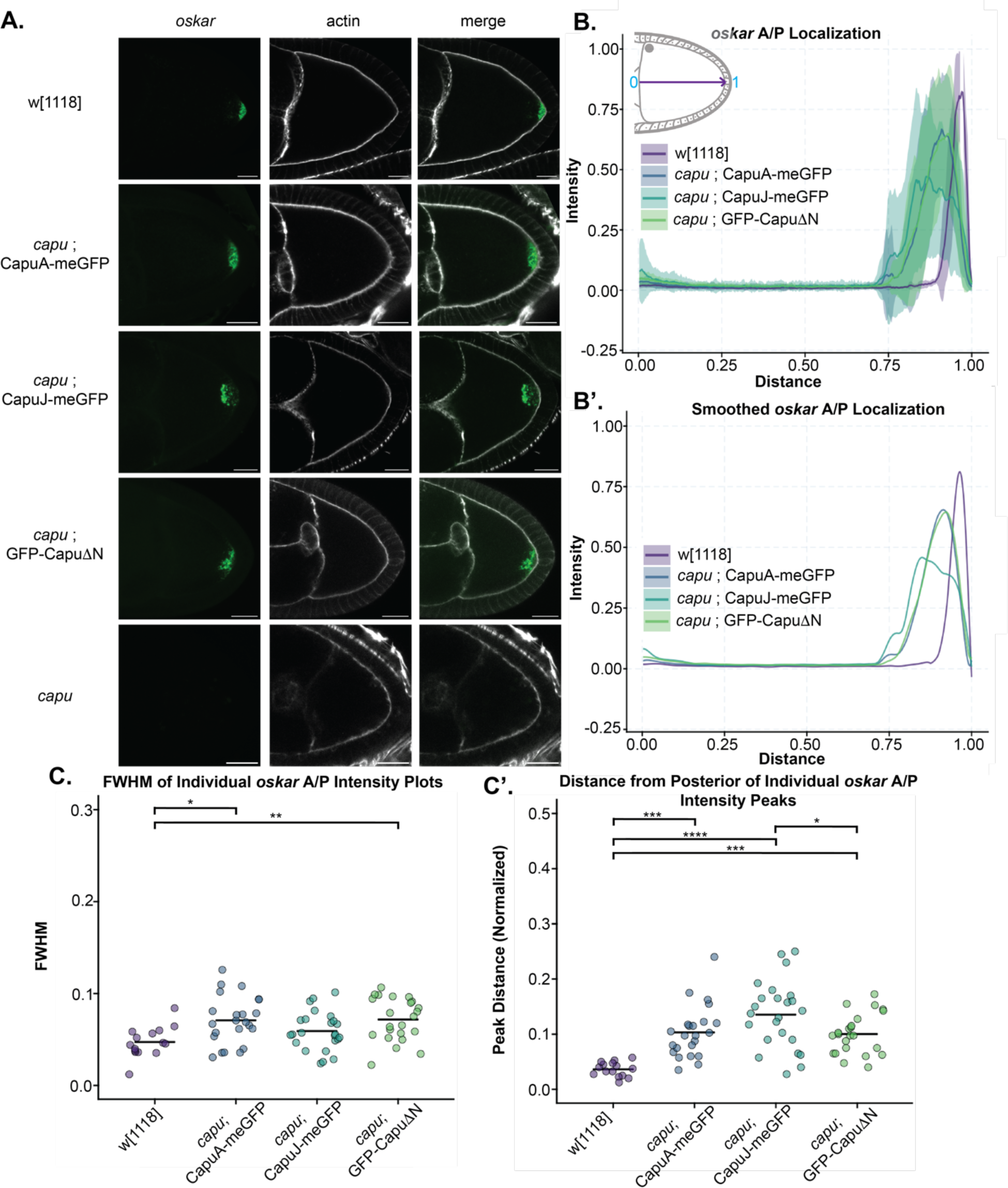
Expression of membrane bound Capu alters posterior mRNA localization. (A) *oskar* mRNA localization was evaluated in stage 9 oocytes using smiFISH (Lu et. al, 2023). Cy3-labeled FLAP-X smiFISH probes recognized the 3’UTR of *osk* mRNA. To determine the localization within the oocyte, AlexaFluor647 Phalloidin was used as a counterstain. Scale bars: 20 µm. (B) Quantification of intensity line scans from the anterior to posterior oocyte for all genotypes overlaid. The anterior/posterior distance was from the nurse cell-oocyte interface to the posterior oocyte cortex-anterior follicle cell border. (B’) Smoothed averages of individual genotype scans; a Savitzky-Golay filter was applied. (C) The full width of the peak intensity at half max of *osk* from individual CapuJ rescue oocytes is not significantly different from wildtype. (C’) Analysis of individual peaks, the distance from the posterior oocyte shifted significantly with CapuJ rescue in comparison to wildtype (****, p = 2.025E-08). Wildtype (w[1118]) n= 15, CapuA rescue n = 23, CapuJ rescue n = 25, CapuΔN rescue n = 23. *capu* null oocytes were not analyzed for localization pattern due to the lack of signal within the oocytes at stage 9.

When *osk* localization is disrupted, translation may not occur (Rongo et al., 1995). As the *osk* peak intensity was so variable with the CapuJ rescue we asked if *osk* was translated. If protein were absent in ∼50% of egg chambers, we would have an explanation for infertility in the CapuJ-GFP background. However, immunofluorescence showed no failure to produce Oskar within the CapuJ-GFP rescue (Fig. S5). Oskar localization patterns were similar to mRNA localization patterns during stage 9 (Fig. 3). When fast streaming begins Oskar that is not anchored can be swept away but there was no apparent difference in retention of Oskar protein at the posterior.

MyoV is the major actin motor involved in mRNA localization at the posterior. Its direct competition with Kinesin-1 within the oocyte is important for transport, eventually taking over in the directed transport of mRNAs into the posterior and anchoring the transcripts, particularly *osk,* during mid-oogenesis (Krauss et al., 2009; Lu et al., 2020). Motor activity of MyoV is directional, migrating towards the growing barbed end of actin filaments and as a consequence is influenced by the orientation of actin filaments within the oocyte. We asked if MyoV localization was altered when Capu remains bound to membranes in the oocyte. We observed a slight disruption of MyoV localization at the posterior in CapuJ-GFP rescues (Fig. 4A). The localization around the cortex is consistent between all genetic backgrounds (Fig. S6B). The A/P localization in CapuJ-GFP oocytes is spread slightly further towards the anterior (Fig. 4B). Measurement of the spread of MyoV signal revealed CapuJ has an approximately 40% greater anterior spread (FWHM = 0.1) compared to wildtype MyoV localization (FWHM = 0.06). CapuA-GFP rescue also exhibited a slight spread of signal towards the anterior on average (FWHM = 0.075) (Fig. 4C’). In contrast, *capu* null oocytes fail to establish a posterior anchor (Fig. S6). Together, these data are consistent with altered *osk* mRNA and protein localization within the stage 9 oocyte and an altered posterior anchor organization due to membrane-bound Capu.

**Figure 4.**
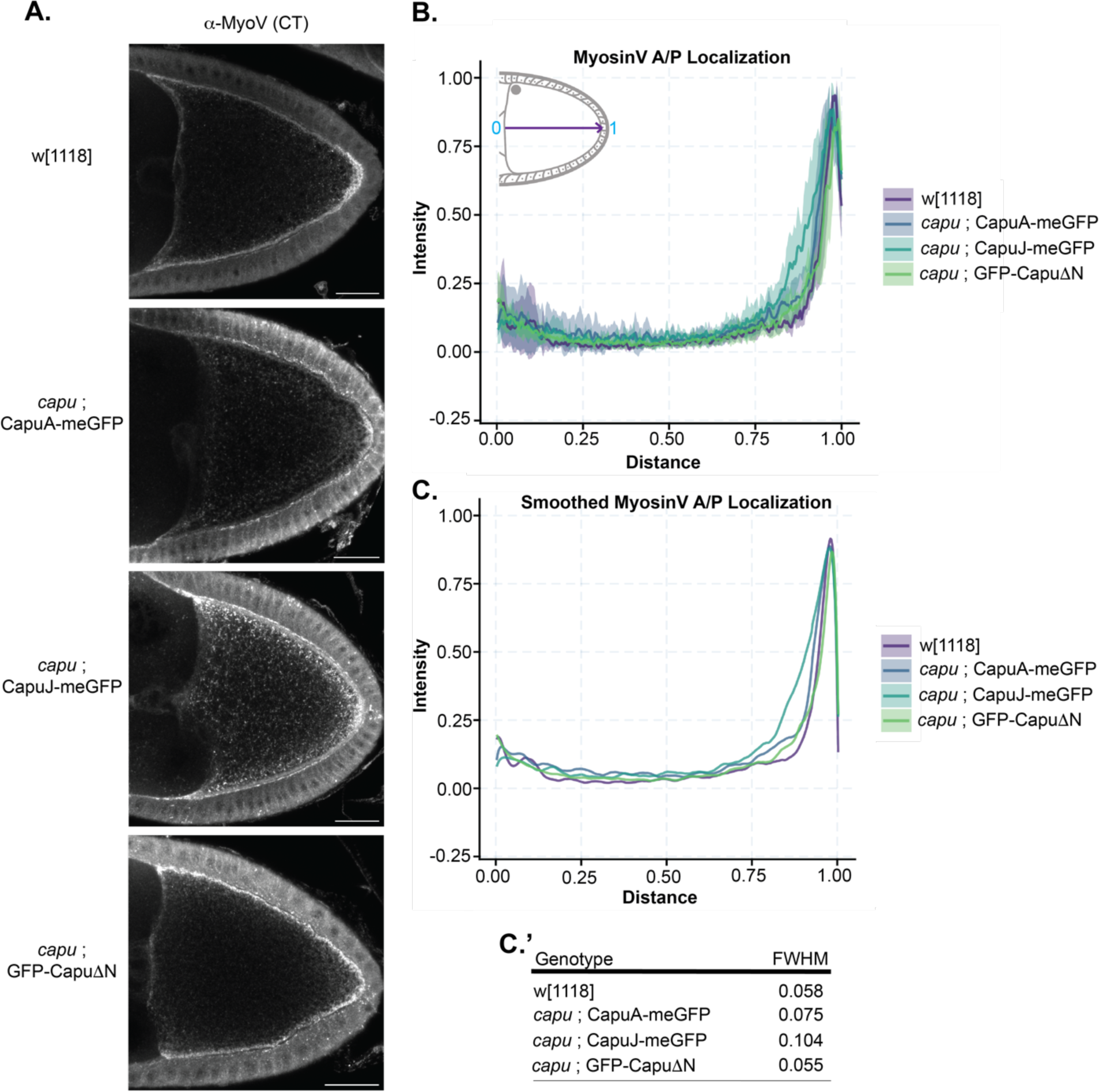
MyosinV at the posterior is slightly altered when expressing membrane bound Capu. (A) MyoV localization was evaluated in stage 9 oocytes using immunofluorescence, with MyoV-CT rabbit antiserum (Gift from the Ephrussi Lab). Scale bars: 20 µm. (B) Quantification of intensity line scans from the anterior to posterior oocyte for all genotypes overlaid. The anterior/posterior distance was from the nurse cell-oocyte interface to the posterior oocyte cortex-anterior follicle cell border. (C) Smoothed averages of individual genotype scans; a Savitzky-Golay filter was applied. (C’) Full width of individual curves at half max (FWHM) were determined. Wildtype (w[1118]) n= 6, CapuA rescue n = 10, CapuJ rescue n = 10, CapuΔN rescue n = 8.

### Failed rescue with myristoylated Capu is not a classic posterior-group phenotype

Given the disruption to *osk* anchoring but apparent recovery of Oskar localization, we asked if posterior patterning was functionally rescued by examining cuticles of offspring from the rescue lines. While CapuA-GFP cuticles were indistinguishable from those of wildtype, the preparations for CapuJ-GFP and GFP-CapuΔN embryos failed, repeatedly. They seemed to lack larvae inside their eggshells that were sufficient to withstand cuticle preparation. Preliminary examination suggests that many of the offspring did not complete embryogenesis.

As an alternate approach, we examined germ cell formation. Germ cell precursors are formed at the oocyte posterior and the process is highly sensitive to disruption of posterior axis determinants; proper localization of Oskar as well as mRNA binding proteins, Vasa, Tudor, and Aubergine are required (Santos and Lehmann, 2004). Penetrance of this phenotype is often evident even if fertility of the parental line and/or the body axes of the offspring are not strongly impacted. That is, female flies with mutations in classical posterior-group genes are grandchildless. We, therefore, asked whether the disruption in posterior organization we observed was hindering pole cell formation. The F1 offspring were crossed to wildtype males and their ovaries were examined as a bulk readout of pole cell formation. Wildtype ovary pairs were less frequently observed in the offspring of both CapuA-GFP and CapuJ-GFP rescues (averages of 68% and 84%, respectively) (Fig. 5). However, given that the CapuJ-GFP offspring had wild type ovaries more often than the CapuA-GFP offspring, we conclude that fertility loss does not reflect a classical posterior-group phenotype. That is, while rescuing oogenesis with membrane bound CapuJ-GFP led to observed differences in the posterior pole, the flies that matured to adulthood were not “escapers” that exhibit signs of posterior defects; body patterning was correct and pole cells still formed.

**Figure 5.**
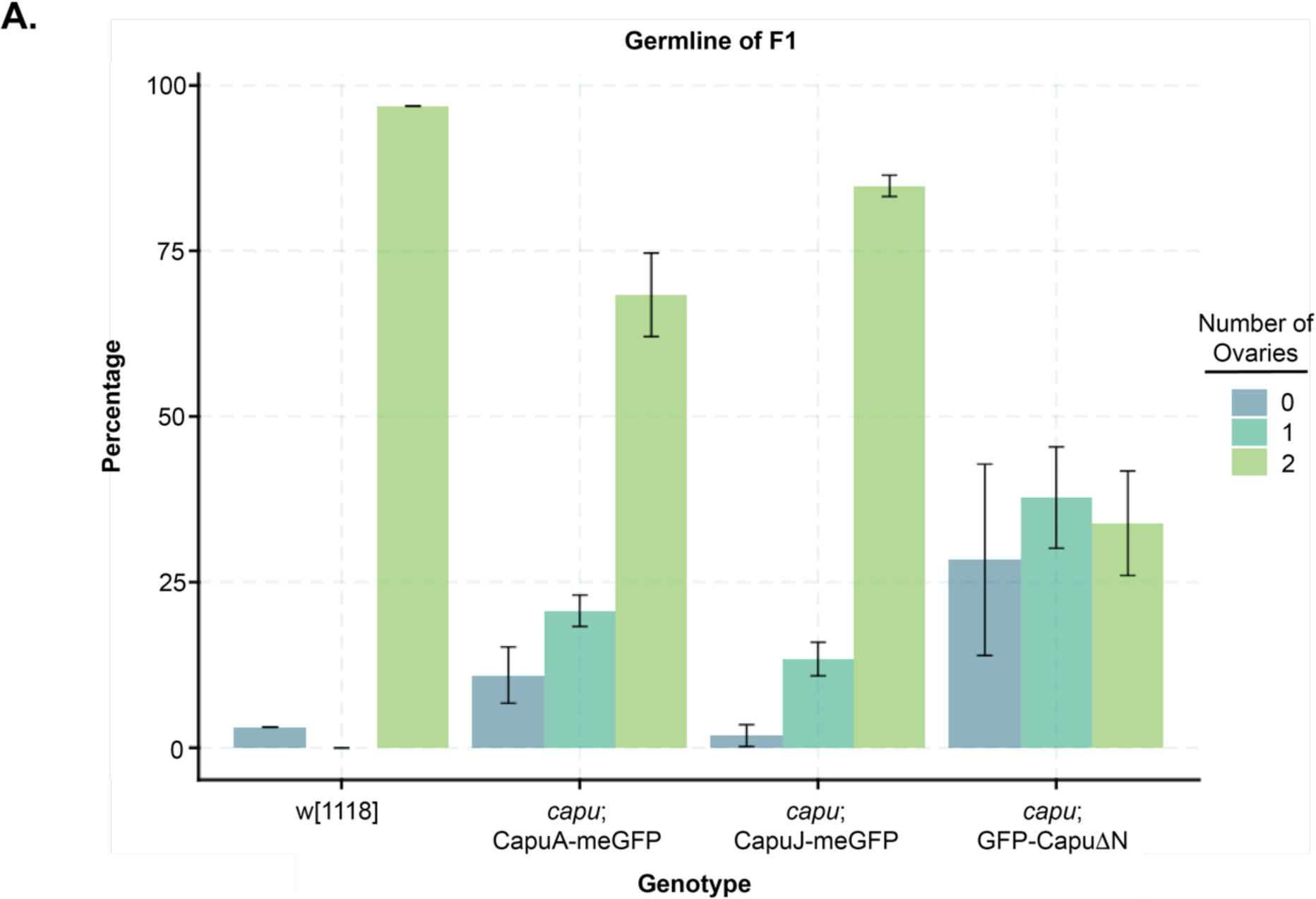
Membrane bound Capu rescue does not exhibit characteristics of the posterior group phenotype. Progeny derived from CapuA or CapuJ rescue females do not show a significant decrease in germline development. Progeny females (F1) from the CapuA (Df(2L)ed^1^/*capu-*Gal4-K10; UASp::CapuA-meGFP/+) and CapuJ (Df(2L)ed^1^/*capu-*Gal4-K10; UASp::CapuJ-meGFP/+) rescue were crossed to wildtype (w[1118]) males and the offspring were allowed to develop. Dissected ovaries were classified as wildtype (two healthy ovary pairs, 2/light green), one ovary (where one ovary appeared wildtype in size and the other shrunken, 1/teal), or none (where when dissected there is somatic tissue but no germ cells are housed in the tissue, 0/blue). Wildtype (w[1118]) n= 32, CapuA rescue n = 130, CapuJ rescue n = 107, CapuΔN rescue n = 116.

The phenotype of GFP-CapuΔN rescue was more consistent with that of a traditional grandchildrenless phenotype. For the GFP-CapuΔN rescue, only 33% of offspring had wildtype ovaries (Fig. 5). Even more had only one ovary. Thus, we conclude that GFP-CapuΔN failed to rescue fertility by a different mechanism than GFP-CapuJ. Apparently, hyperactivity, due to loss of autoinhibition, is more detrimental than mislocalization of Capu.

### Is the function of Capu at the posterior independent of Spir?

Spir and Capu function together to build the actin mesh during mid-oogenesis (Dahlgaard et al., 2007; Quinlan, 2013). Given that CapuJ-GFP localization is so different from CapuA-GFP but still builds a functional mesh, we asked if it impacts Spir localization. To better determine where Spir is localized, we used Crispr/Cas9 to add either smGFP-HA (Nern et al., 2015) or mScarlet-HA at the C-terminus. Fertility rates of females expressing homozygous Spir edits were 88% and 94%, respectively. As observed with immunofluorescence and similar to transgene localization, Spir is enriched at the oocyte cortex (Fig. S7A). Furthermore, we were able to confidently identify Spir-positive punctae which we presume to be vesicles. These punctae are found throughout the oocyte but at higher concentrations near the cortex (Fig. S7B). Like Capu, we observed Spir expression in the border cells. We also observed Spir in the follicle cells surrounding the oocyte (Fig. S7A,B). Interestingly, the concentration of Spir at the posterior of the oocyte is lower than other places. The decrease in posterior Spir was consistent in both endogenously tagged lines (Fig. 6, Fig. S7C). It becomes very apparent when comparing Spir and Capu localization (Fig. 6). Comparison of rescue with CapuA-GFP and CapuJ-GFP shows no detectable difference in Spir localization pattern (Fig. 6), suggesting that Capu is not determining Spir localization. Decreased Spir at the posterior cortex and the fact that Spir localization is not impacted by the Capu isoform could indicate that Capu works independently of Spir at this location. Of note, now that we know that Spir is expressed in the follicle cells, in addition to the oocyte, further work is required to determine if this decrease is in one or both of these cell types.

**Figure 6.**
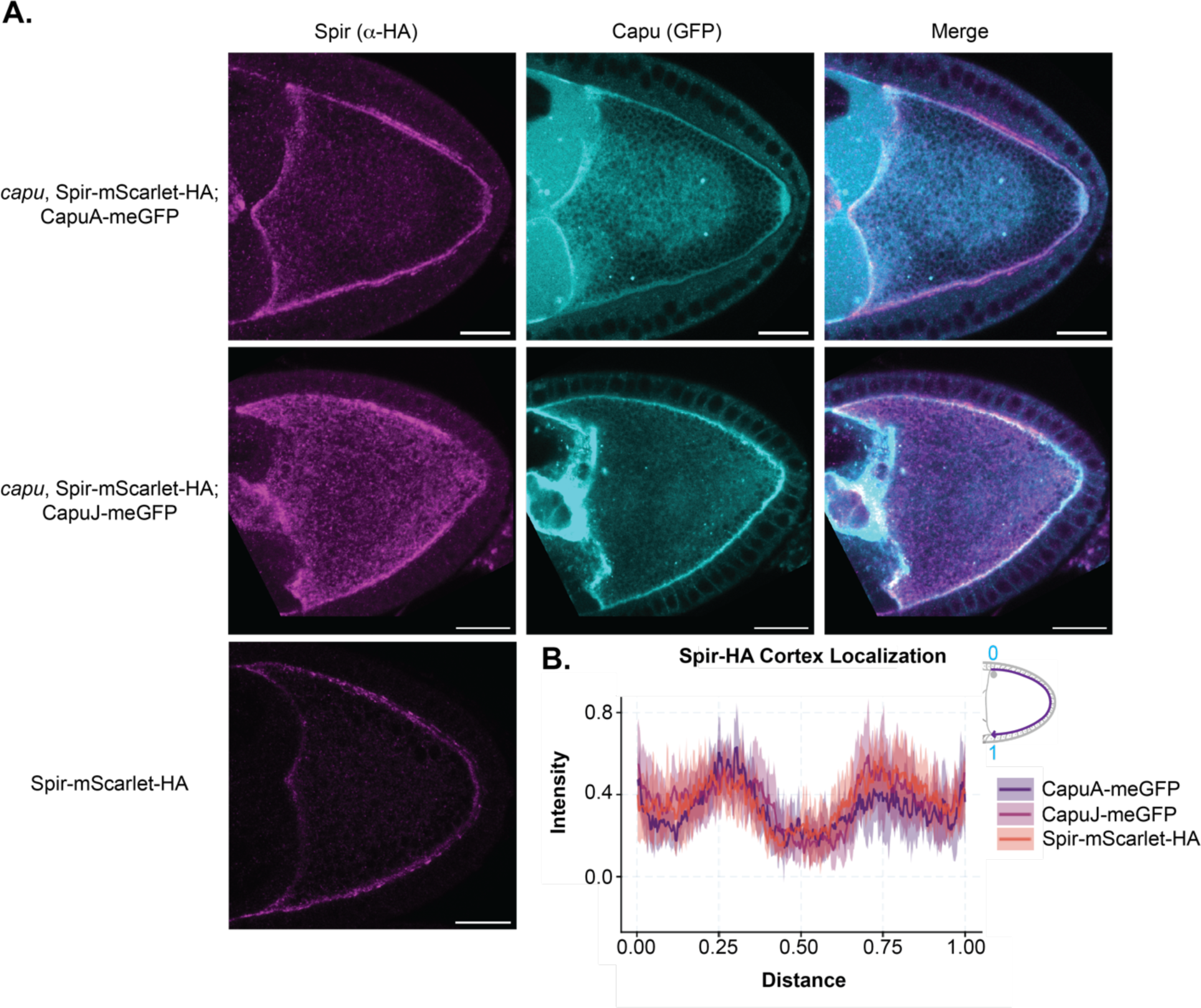
The posterior function of Cappuccino may be independent of Spire. (A) Co-imaging of stage 9 egg chambers expressing Capu-GFP (direct fluorescence of the transgene) or wildtype Capu and Spir-mScarlet-HA (endogenously tagged, stained with α-HA). There is a striking lack of Spir at the posterior end of the oocyte. Scale bars: 20 µm. (B) Quantification of Spir signal around the cortex of the oocyte. A dip in intensity is apparent at the posterior in all backgrounds. Wildtype Capu background (Spir-mScarlet-HA) n= 9, CapuA rescue n = 7, CapuJ rescue n = 8.

## Discussion

In the developing mouse and fly oocytes, Spir and Capu (FMN2) fill the cytoplasm with an actin mesh (Azoury et al., 2008; Dahlgaard et al., 2007; Schuh and Ellenberg, 2008). FMN2 colocalizes with Spir on vesicles in the mouse oocyte (Pfender et al., 2011). We now confirm that Capu is not enriched with Spir on vesicles in the *Drosophila* oocyte. As suggested by transgene expression, endogenously edited genes reveal diffuse localization of Capu throughout the *Drosophila* oocyte (Fig. S1). We also detected relatively strong expression of Capu in specialized somatic cells, including the border cells and polar cells (Fig. S1). We did not pursue this observation, however, because our rescue experiments demonstrated that germline expression is sufficient for oogenesis (Table 1, CapuA rescue).

Localization of Spir and Capu is independent of one another within the *Drosophila* oocyte, as demonstrated by the lack of colocalization and the unaltered localization of Spir when CapuJ is expressed and enriched at membranes (Fig. 6). We would not have predicted this independence based on their colocalization in the mouse oocyte and our finding that Spir is sufficient to drive Capu localization in S2 cells (Pfender et al., 2011; Vizcarra et al., 2011). Further, genetics demonstrate that the pair of actin nucleators must physically interact to function in *Drosophila* oogenesis (Quinlan, 2013). Together, these observations raise the questions of when, where, and for how long Spir and Capu interact? The data suggest that the interactions are transient and that Capu elongates barbed ends of filaments away from Spir-enriched surfaces (Bradley et al., 2019), which would likely impact the organization and function of the structures built. Given that Spir and Capu bind with nanomolar affinity in vitro (Vizcarra et al., 2011), the data also suggest that the interaction is somehow down regulated in the oocyte. It follows that this regulatory mechanism is absent in S2 cells. Finally, the distinct localizations suggest that Spir and Capu may not always function as a team. If so, do one or both of these nucleators build structures other than the mesh? It has been difficult to address this question, as premature fast steaming due to absence of the actin mesh has such drastic consequences on development.

We were particularly interested in the marked decrease of Spir at the posterior of the oocyte, despite no change in Capu along the cortex (Fig. 6). Further, if CapuJ was the isoform of Capu expressed, we observed displacement of *osk* mRNA during mid oogenesis (Fig. 3). In fact, in all three rescue backgrounds, CapuA, CapuJ, and CapuΔN, the spread of *osk* around the cortex was broader than observed in wildtype oocytes (Fig. S4). We propose that the increased region or *osk* retention reflects a role for Capu in posterior anchoring. That is, the anchoring site is expanded due to high levels of expression of any of the Capu the transgenes. However, altered A/P localization of *osk* was distinct to CapuJ expression (Fig. 3A). Mislocalization of *osk* at the posterior has been investigated in a number of genetic backgrounds. Proper organization of the cytoskeleton and motor activity is required to transport and anchor *osk* in a posterior cap. Mutants of par-1, khc, didum among others often result in *osk* at a central dot, reflecting failure to transport *osk* along properly organized microtubules and capture it with MyoV at the posterior (Cha et al., 2002; Doerflinger et al., 2006; Doerflinger et al., 2022; Krauss et al., 2009; Lu et al., 2020; Parton et al., 2011; Zimyanin et al., 2008). In the absence of Long Osk, *osk* spreads continuously from the posterior, reflecting failure to anchor (Vanzo and Ephrussi, 2002). The CapuJ phenotype is distinct from both of these. While the average location of *osk* was spread broadly during stage 9, within any individual oocyte, the *osk* was tightly gathered (Fig. 3C). We interpret this as evidence that altering Capu geometry (by myristoylation) reveals a role for Capu in localizing but not building the actual “posterior landing platform”. We observe *osk* translation both at the posterior and at what we are referring to as the displaced posterior landing platform (Fig. S5). The fact that *osk* is translated indicates that functional organization of posterior determinants is accomplished. Development of germ cells and deviation from typical posterior group phenotypes further strengthens this interpretation. In the future, it will be important to identify the minimal elements of the landing platform.

Finally, if posterior determination is slightly perturbed but ultimately functional, why is fertility decreased when CapuJ is expressed? There could be additional structures built by Capu that are defective. There are previous reports of long posterior actin filaments that are induced by Long Oskar and Capu during mid-oogenesis that further anchor Short Oskar for continual delivery of posterior pole plasm components (Chang et al., 2011; Tanaka et al., 2011). We would assume that by greatly increasing the localization of Capu to the posterior oocyte (Fig. S1) that we would see a dramatic increase in filament formation. We were unable to observe such a phenomenon. Perhaps, this is due to technical challenges in distinguishing these filaments from the mesh. We also note that a recent publication suggests the previously described posterior actin filaments are projections, filopodia from the surrounding follicle cells (Mallart et al., 2024). Alternatively, decreased fertility could imply that the mesh has another role that cannot be effectively performed when built with CapuJ-GFP. This role would be in addition to limiting microtubule movement and regulating the transition from slow to fast streaming.

## Materials and Methods

### *Drosophila* Line Generation

#### CRISPR: Spir-mScarlet-HA

Spir was tagged with mScarlet-HA (mScarlet-HA) at its endogenous locus using CRISPR/Cas9 mediated homologous recombination. To generate tagged flies, we followed a combined approach using dual gRNA sequences(Bence et al., 2017)and short homology arms (Kanca et al., 2019). gRNA sequences for the C-termini of Spir (5’ GTCGGCCCTGGATCTGACGCCCGTC 3’) and following the 3’UTR (5’ GTCGGCAAACTAAAGAACAAGATTC 3’) were selected using the CRISPR Optimal Target finder (http://targetfinder.flycrispr.neuro.brown.edu/index.php). Oligonucleotides were cloned into pCFD3-dU6:3gRNA (Addgene #49410) and cloned using the previously designed strategy (Port et al., 2014). 200nt homology sequences with the PAM removed were cloned into homologous recombination vector, a self-linearizable pUC57 (Kanca et al., 2019). The repair template includes a C-terminal mScarlet-HA, the endogenous 3’UTR of Spir, and the fluorescent eye reporter 3xP3_dsRed flanked by two PiggyBac recombinase sites. Plasmids expressing the guide RNAs and donor template were mixed and co-injected in embryos *nos-Cas9* embryos (BDSC 78782,(Ren et al., 2013)) at concentrations of 100ng/μL and 250ng/μL respectively. Offspring were screened via fluorescence of the 3xP3_dsRed for integration of the repair within the genome. Clones were balanced on chromosome II and Cas9 expressing chromosome removed. All injections and initial screening were completed by BestGene (Chino Hills, CA). Proper insertion of mScarlet-HA was confirmed via genomic PCR (forward primer: 5’GGGGATTCAACCTGTTCTCCT 3’, reverse primer: 5’TGTGCAAGTGCGTTCTGAAG 3’) and western blot prior to further experimentation.

#### UASp::CapuJ-meGFP-K10 Generation

CapuJ-meGFP transgene was generated by modifying our CapuA-meGFP expression vector. The coding sequence of CapuA-meGFP was amplified from pTIGER (Quinlan, 2013) lacking the first 238bp (N-terminal exon) to generate a generic Capu-meGFP plasmid in pDONR221. A gene fragment containing the first 326bp of CapuJ (CG93399-RJ) were cloned into the N-termini at a XhoI site. The insert was cloned into pPW (DGRC 1130) modified to include an attB1 site, pPW-attB1, via LR reaction. The final plasmid was then integrated at the attP2 landing site by BestGene.

#### capu-Gal4-K10 Generation

*capu-*Gal4-K10 driver was generated by first adding the K10 3’ UTR terminator site to replace the HSP70 3’UTR within the Trojan-Gal4 plasmid, pBS-KS-atB2-SA(0)-T2A-Gal4 (Addgene, Plasmid #62899). This plasmid was then sent for injection to be integrated in the MiMIC landing site of Capu, MI057537.

Contact the corresponding author for plasmids or fly lines described here.

### Other *Drosophila* Stocks used in this study

UASp::CapuA-meGFP-K10 (Quinlan, 2013), UASp::GFP-Capu-ΔN (Gift from St. Johnson lab (Dahlgaard et al., 2007)), w[1118] (Bloomington, BDSC 3605), Df(2L)ed1/CyO (Bloomington, BDSC 5330)

### Fertility Assays

Approximately 100 test females were crossed to 50 wild-type (wt) males and raised on apple plates for 2 nights at 25°C. On day 3, to synchronize egg laying, flies were pre-cleared on a fresh apple plate containing yeast paste for 1.5 hours. A fresh plate was added, also containing yeast paste, and eggs laid over the following 3-hour time period were collected. Approximately 200 eggs were laid during that window. Eggs were moved to a new apple plate using a paint brush and incubated at 25°C for 24 hours. The number of eggs that hatched during that window were recorded. Each trial was repeated with independent crosses three times. The data reported in the table is an average across the three trials to obtain the fertility rate.

### F1 Ovary Dissection/Germline Perdurance

Approximately 15 F1 females, progeny of the test flies, were crossed to 7 wild-type males. Female progeny from these crosses were then collected, aged to 3 days post-eclosion, and fattened overnight at 25°C. Their ovaries were then dissected in 1XPBS and classified in three categories: wildtype (two average size ovaries), 1 ovary, (one ovary with one shrunken ovary), or 0 ovaries (somatic tissue/both shrunken). Each trial was repeated with independent crosses three times. The data reported in the graph is an average across the trials, error bars are +/- one standard deviation.

### Microscopy and staining

All microscopy images, live and fixed, were collected on a Zeiss LSM 780 confocal microscope. Flies were raised at 25°C and fed yeast paste for 16-24 hours prior to imaging.

The actin mesh was stained as described (adapted from (Dahlgaard et al., 2007; Quinlan, 2013)) using 1μM AlexaFluor488-phalloidin (Thermo Fisher Scientific, A12379) or 1μM AlexaFluor647-phalloidin (Thermo Fisher Scientific, A22287) for 45 minutes.

Immunofluorescent staining of the HA (Spir-mScarlet-HA) and MyoV was performed by dissecting ovaries in cold 1XPBS, and fixing in 5% PFA (Electron Microscopy Solutions, 15714), diluted in 0.16X PBS with Heptane, protocol is modified from (Robinson and Cooley, 1997). Samples were stained with 1:1,000 rabbit anti-HA (C29F4) (Cell Signaling Technology (CST), 3724S), 1:3,000 rabbit anti-Osk NT (gift from A. Ephrussi (Bose et al., 2022)), or 1:500 rabbit anti-MyoV CT (gift from A. Ephrussi (Krauss et al., 2009), preabsorbed with fixed and unblocked wildtype ovaries) respectively overnight at 4°C. 1:200 secondary was used, AlexaFluor568-Donkey α Rabbit (Invitrogen, A10042). Sometimes with AlexaFluor647-phalloidin counterstain (Thermo Fisher Scientific, A22287).

Live egg chambers were dissected under halocarbon 700 oil (Sigma-Aldrich, H8898). For GFP-fusion localization imaging, egg chambers were excited with 488 nm laser using 40X oil immersion objective. Images were captured at 1024x1024 resolution with 4 frame averaging. Z-stacks were collected with 1 μm steps over a range of 5-10 μm. Autofluorescent yolk granules were excited using a UV 405nm laser. To track fluid flows, images were captured every 5 seconds for a 5-minute duration. Maximum ‘time’-projections were created in Fiji for representation of motion throughout the movie for the oocyte.

### Analysis of Streaming: Particle Image Velocimetry and Correlation Functions

Cytoplasmic streaming velocities were determined from confocal images using a particle image velocimetry lab (PIV lab) MATLAB package generated by William Thielicke (Thielicke and Sonntag, 2021; Thielicke and Stamhuis, 2014).

ROIs were drawn around the oocyte and the background masked so only the area containing yolk (the oocyte cytoplasm) was interrogated. PIV algorithm used was FFT window deformation, a direct Fourier transformation correlation with multiple passes and deforming windows. The passes were interrogation areas of 30px with a 15px step (50%), decreasing to 20px and 10px. This allowed for interrogation areas that were specific to yolk granule size and changing with pattern to be analyzed for each frame. Limits were selected to refine data, and PIV lab was permitted to interpolate missing data based on surrounding values. From this analysis the vector information was extracted for the motion stream and the mean streaming velocity was determined for each oocyte analyzed. Mid and late-stage oocytes were analyzed using this method.

Output raw files of vector information were then further processed in custom Correlation analysis software, 2DVelocityCorrelation, which can be found on GitHub (https://github.com/LiamABailey/2DVelocityCorrelation). Work was based on (Dombrowski et al., 2004; Ganguly et al., 2012). For each frame of PIV, a correlation function was determined and then averaged to generate the correlation for the oocyte. For each oocyte we determined the maximal radius at which the vectors were half correlated and plotted in R using ggplot2. Statistics were run in R using rstatixs package, a one-way ANOVA with Tukey-Kramer post-hoc test to determine significance between genotypes. See graphs for information about p-values generated.

### Quantification of Staining (Immunofluorescence and smiFISH)

To determine localization of *osk* mRNA, Spir-mScarlet-HA, and MyoV, in the oocyte, images were analyzed using the line scan tool in Fiji. 5 or 10μm Z-stack data were collected as described above. A maximum intensity projection was made in FIJI. For mRNA localization analysis, the mRNA/Cy3 signal was overlayed with the actin staining to determine bounds of the oocyte. Anterior to posterior oocyte (anterior follicle cell) lines were drawn, as well as around the oocyte cortex - excluding the anterior oocyte-nurse cell border. A line width of 20px was used and intensity for the channel of interest was plotted. Data were saved from this, intensity and distances measured were normalized using python scripts. Data were then further analyzed in python and plotted in R using ggplot2. Averages for each genotype were smoothed using python, SciPy, Savitzky-Golay filter. From this the maximum peak metrics were determined: full width at half max and peak distance from the posterior oocyte.

### Single-Molecule Inexpensive Fluorescence in situ Hybridization (smiFISH)

Our protocol was based on (Calvo et al., 2021; Lu et al., 2023; Tsanov et al., 2016) and generously provided by Lu/Gelfand. Twenty-base-long DNA probes with complementarity to the mRNA *bcd* 3’UTR*, grk* whole mRNA sequence, and *osk* 3’UTR, with 3’ FLAP-X complementary probe (5’-CCTCCTAAGTTTCGAGCTGGACTCAGTG-3′) were generated using LGC Biosearch Technologies’ Stellaris RNA FISH Probe Designer (masking level five, minimal spacing two bases). 25 probes for each mRNA were ordered from Integrated DNA Technologies (IDT) (probe sequences are listed in Supplement, Table S1). Cy3-FlapX probe was also synthesized by IDT. Ovaries were hybridized in 2μL annealed probe diluted in prewarmed smiFISH Hybridization buffer overnight (>16h) at 37°C in the dark. Egg chambers were counterstained with 0.2μM AlexaFluor647-phallodin (Thermo Fisher Scientific, A22287) for 10 minutes and mounted on slides in Prolong Gold antifade with DAPI (Thermo Fisher Scientific, P36931).

---

**Figure S1:**
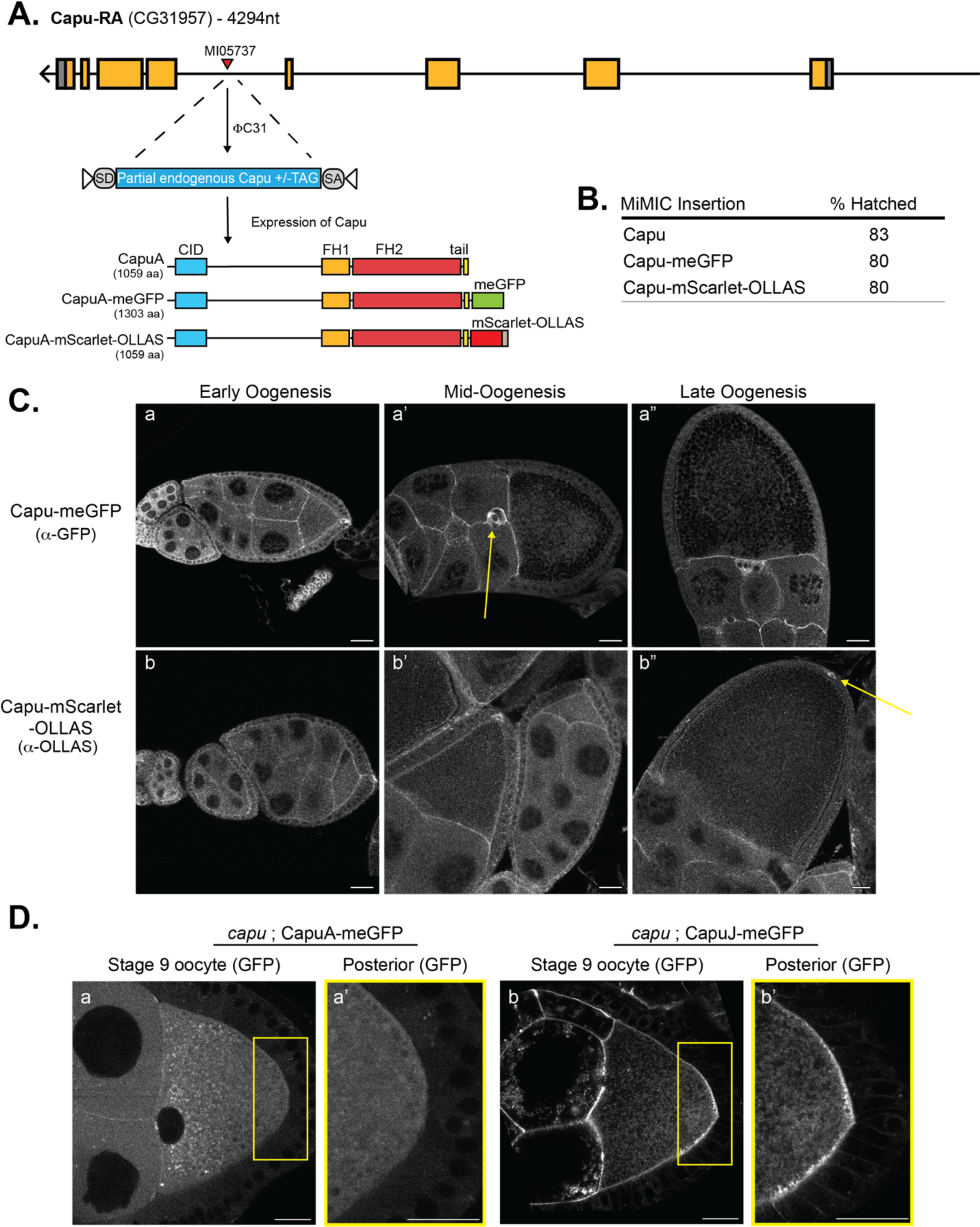
Determining the localization of Capu. (A.) Schematic of insert strategy used to generate endogenously tagged Capu lines. C-termini of Capu-RA is shown as an example, all isoforms of Capu contain MI05737. (B.) Fertility of endogenously tagged Capu lines generated. The % hatched is reported as the average of three independent trials. Fertility is reported as the percentage of eggs that hatched within 24 hours of being laid. All test female flies were crossed to wild-type males to evaluate the maternal contribution to the rescue rate. (C.) Immunofluorescent staining of endogenously tagged Capu in egg chambers throughout oogenesis. Capu-meGFP (a-a”) and Capu-mScarlet-OLLAS (b-b”) staining are consistent with transgene rescue data. Scale bars: 20µm. (D.) CapuA-meGFP (a-a’) and CapuJ-meGFP (b-b’) stage 9 egg chambers. Posterior localization pattern varies between isoforms. CapuJ (b’) is highly enriched at the cortex. Scale bars: 20µm.

**Figure S2:**
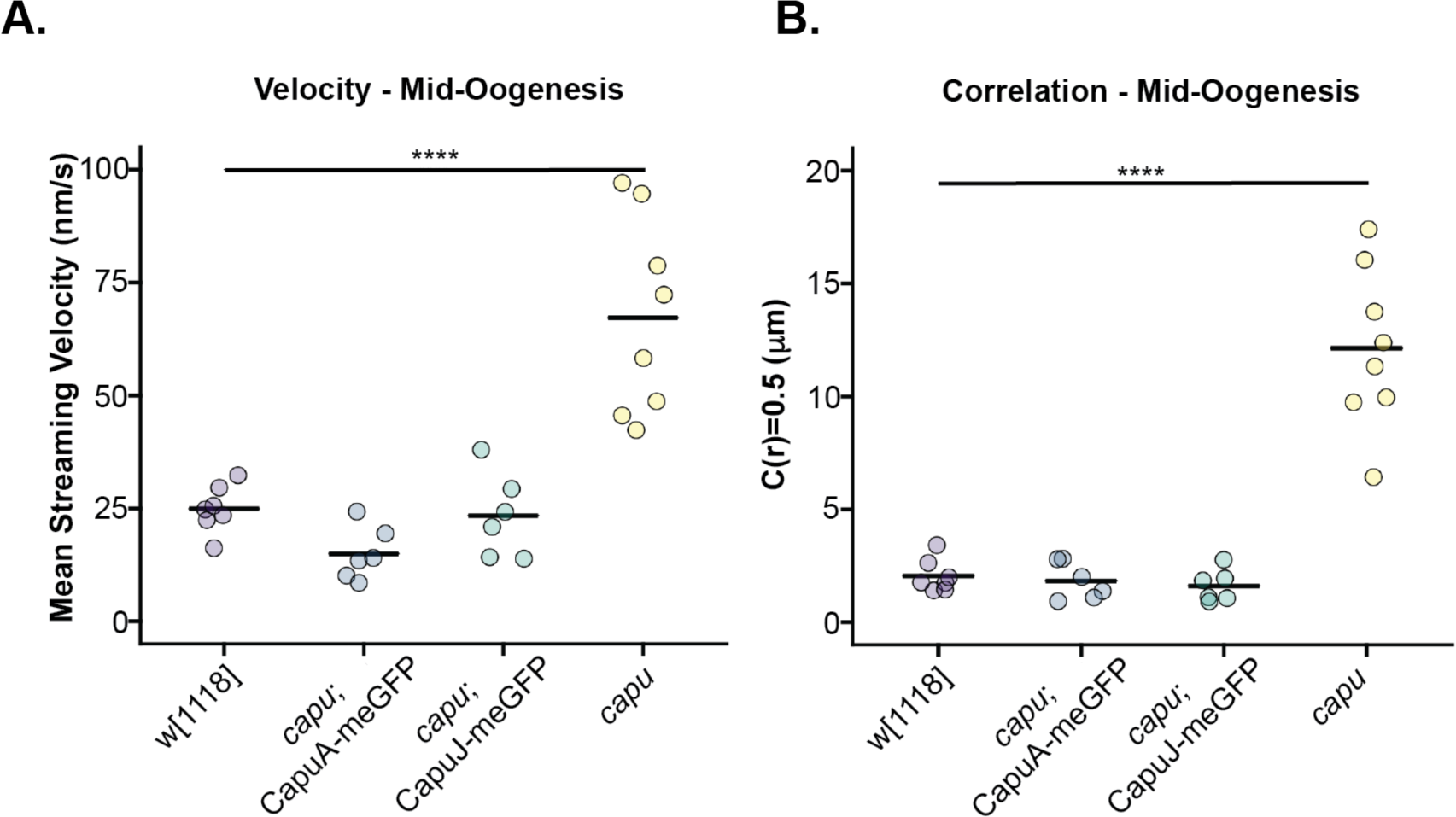
Analysis of streaming velocities and patterns during mid-oogenesis. (A-B) Dot plots of streaming velocities and patterns (correlation radius = 0.5), the bar in each dataset indicates the average value. N ≧ 6 for each genotype analyzed. (A) Quantification of streaming shows no significant diferences between the wildtype and Capu rescue backgrounds during mid oogenesis (∼25nm/s). Premature fast streaming in *capu* null oocytes is accelerated three-fold (∼65nm/s) and is significantly diferent from the other genotypes as indicated (****, p = 0.00001652). (B) Analysis of the mid-oogenesis streaming patterns, correlation (described in the methods), using a custom code (2DCorrelation, LiamABailey, GitHub). No significant diference is detected between wildtype and CapuA or CapuJ rescue flies (∼2µm). *capu* null oocytes exhibit a significant increase in correlation radius, ∼13µm (****, p = 1.4E-08).

**Figure S3.**
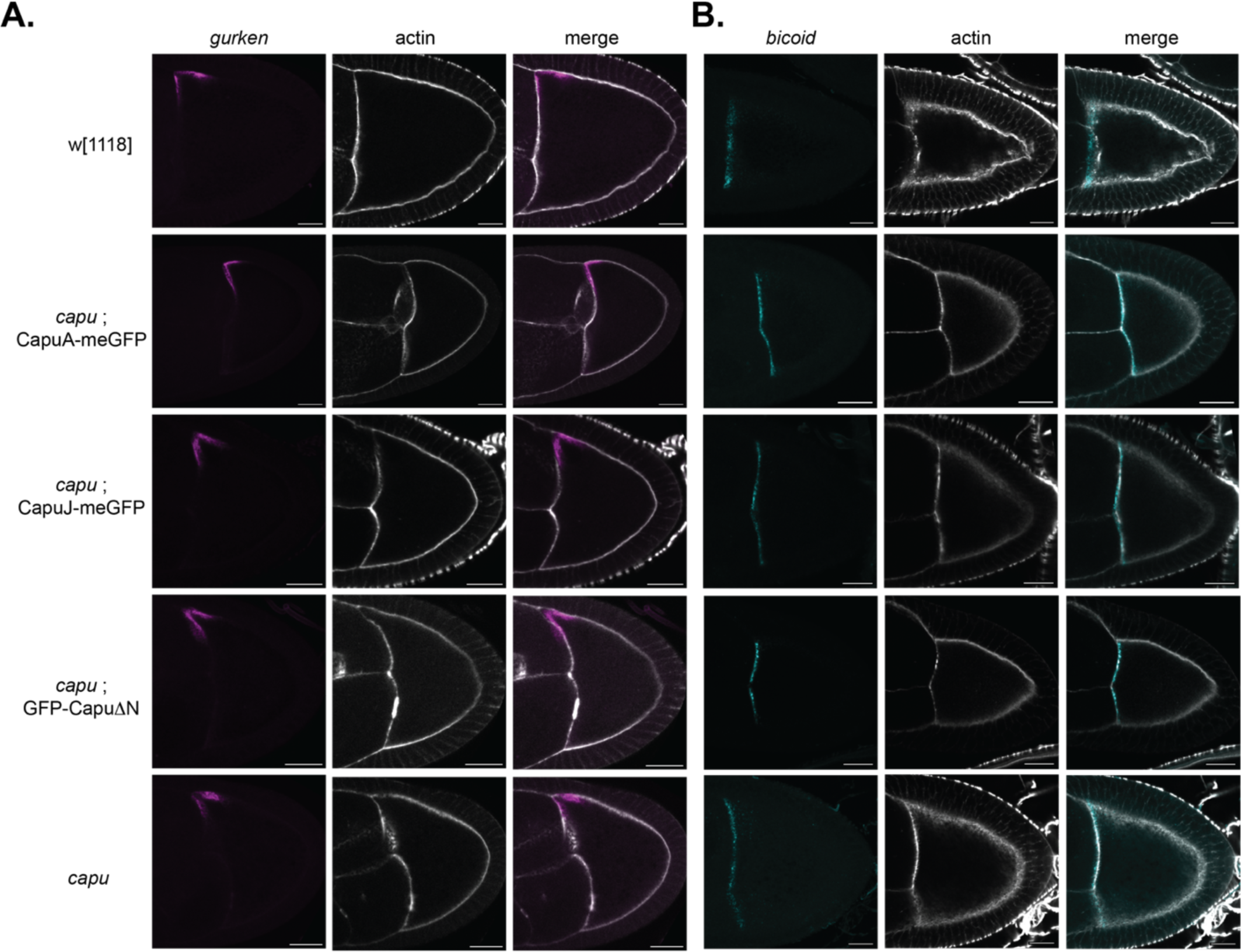
Anterior mRNA localization is unaCected by altering Capu localization within the *Drosophila* egg chamber. (A.) *gurken* mRNA localization was evaluated in stage 9 oocytes using smiFISH (Lu et. al, 2023). Cy3-labeled FLAP-X smiFISH probes recognized the entire mRNA sequence of *grk*. Scale bars: 20µm. (B.) *bicoid* mRNA localization was evaluated in stage 9 oocytes using smiFISH (Lu et. al, 2023). Cy3-labeled FLAP-X smiFISH probes recognized the 3’UTR of *bcd*. Scale bars: 20µm.

**Figure S4.**
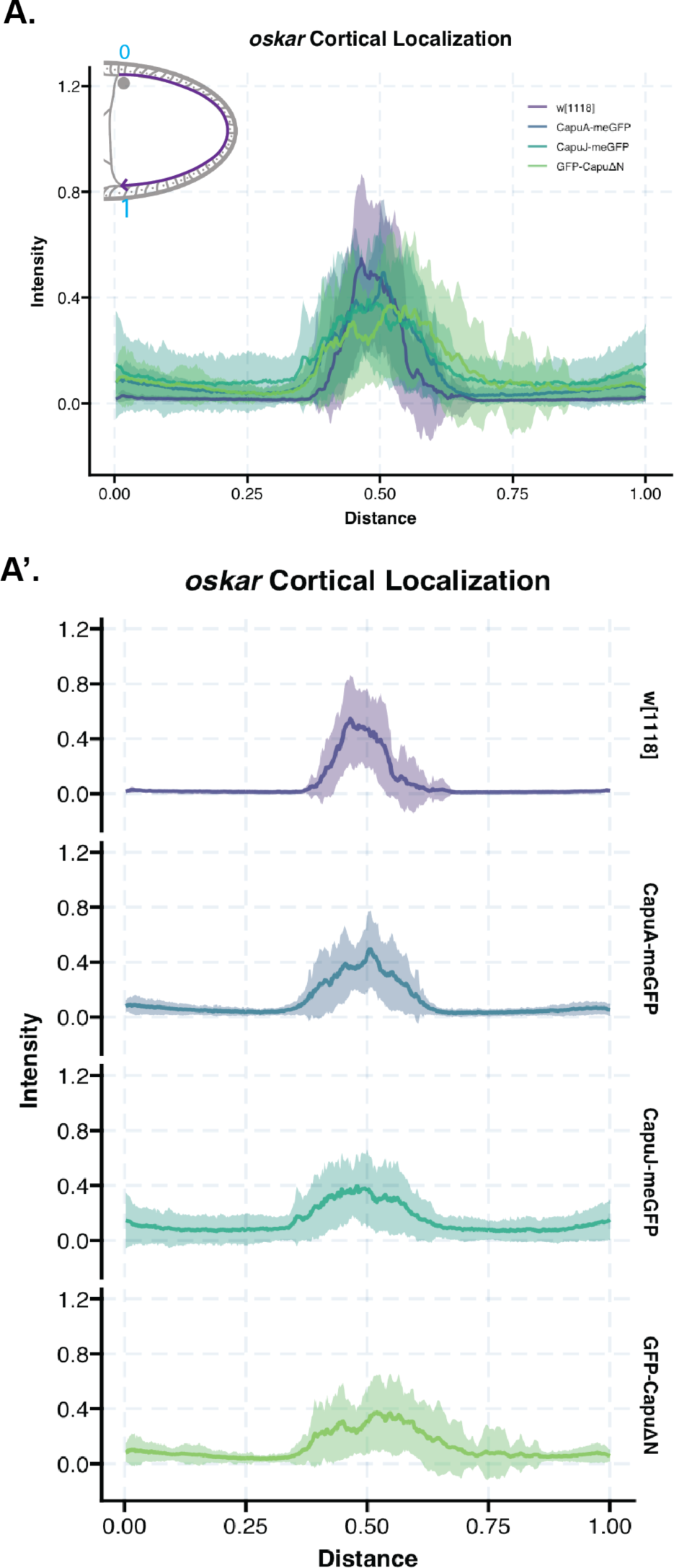
*oskar* mRNA localization around the cortex is expanded but not significantly diCerent between Capu rescue backgrounds. (A.) Quantification of normalized intensity line scans around the oocyte cortex for all genotypes overlaid. Rescue oocytes exhibit a decreased posterior intensity with greater spread, compared to wildtype oocytes. (A’) Individual genotype scans, average (solid line) with the standard deviation. Wildtype (w[1118]) n= 15, CapuA rescue n = 23, CapuJ rescue n = 25, CapuΔN rescue n = 23. *capu* null oocytes were not analyzed for localization pattern due to the lack of signal within the oocytes at stage 9.

**Figure S5.**
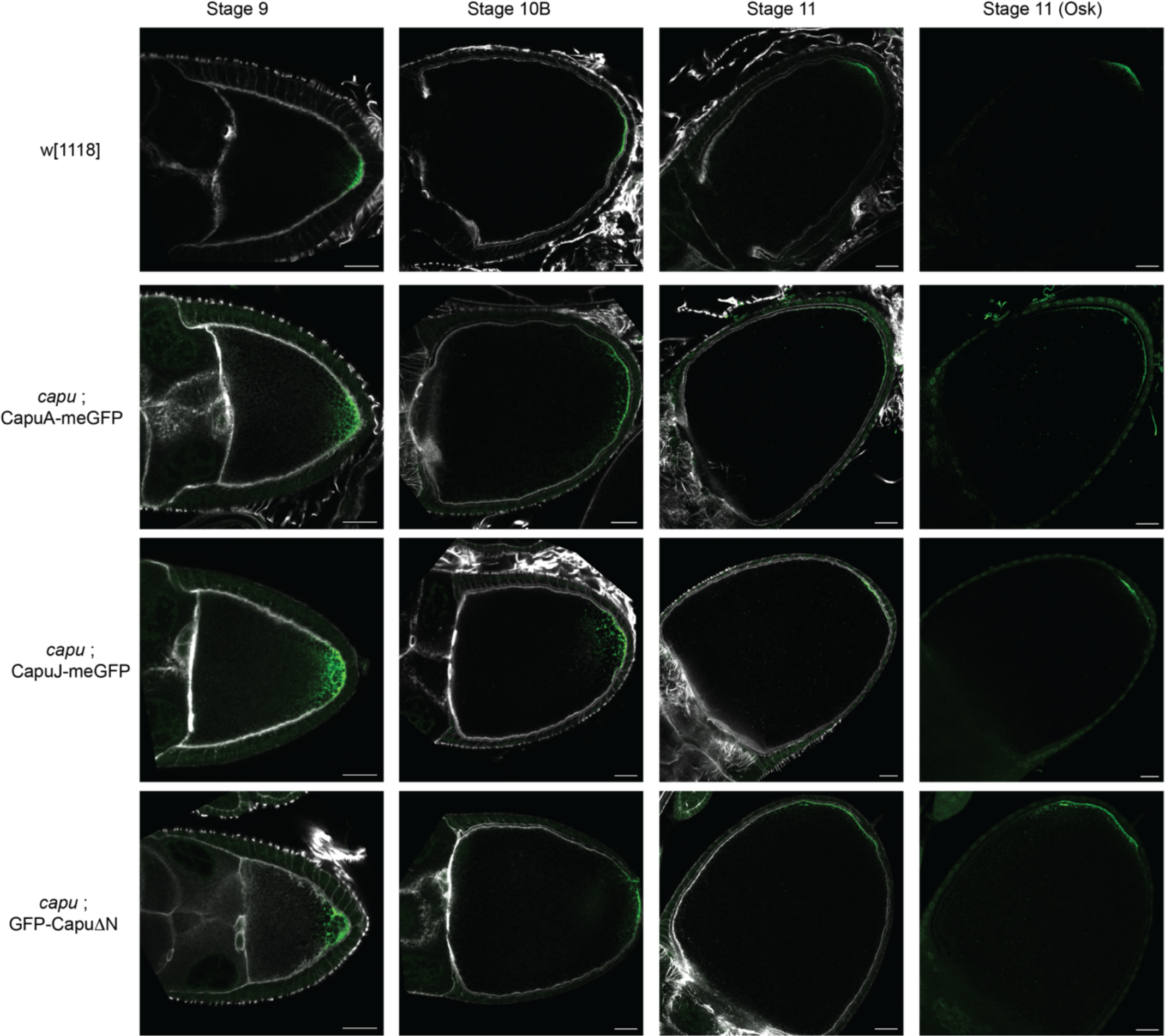
Oskar translation occurs when Capu localization is altered. (A.) Staining for Oskar (1:3000, gift from Ephrussi Lab) at mid (stage 9) and late oogenesis at the onset (stage 10B) and as streaming velocity increases (stage 11). At all stages and in all rescues, Oskar is present. Scale bars: 20µm.

**Figure S6.**
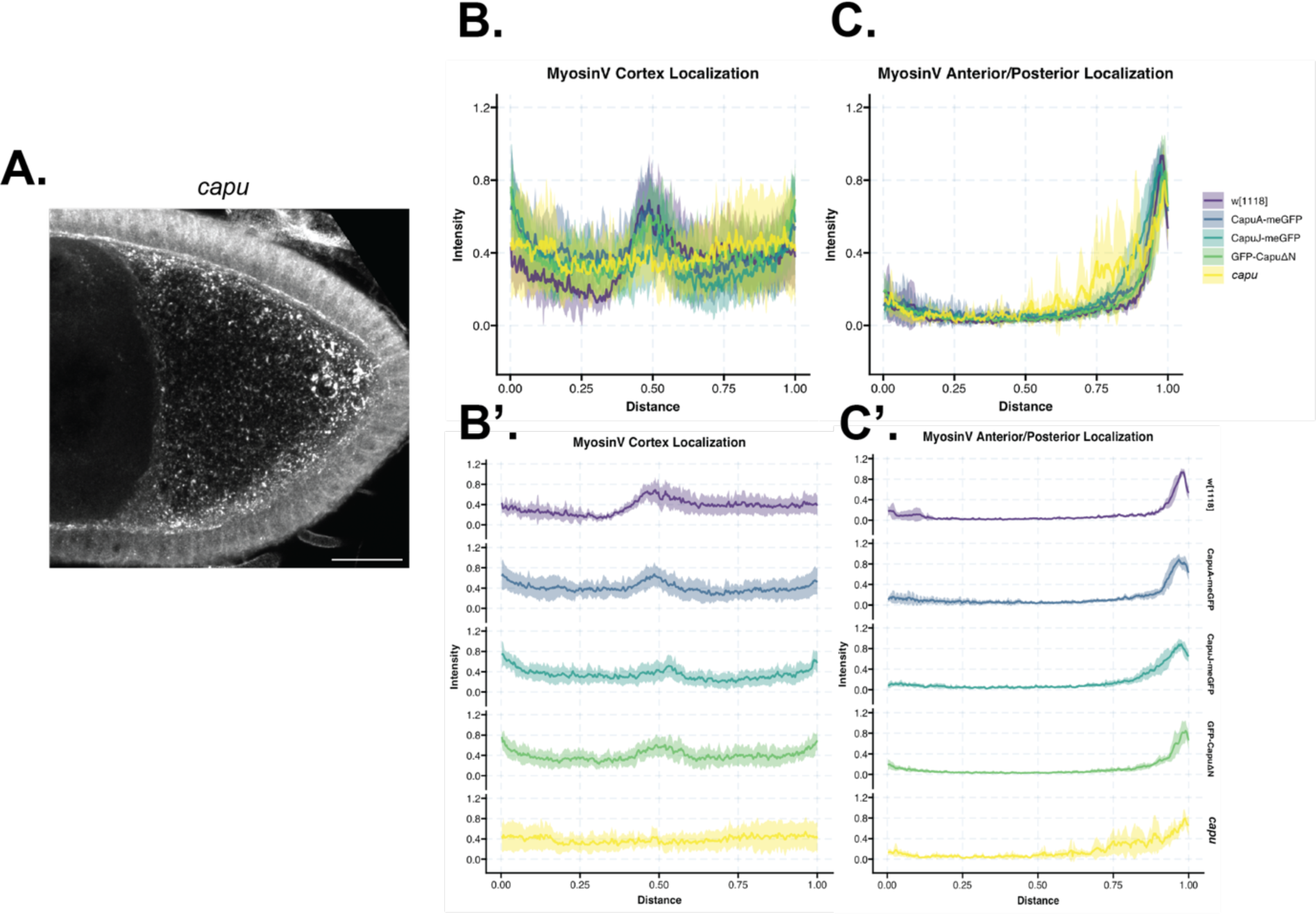
MyoV distribution is significantly disrupted in *capu* null oocytes. (A) MyoV localization in *capu* null stage 9 oocytes using immunofluorescence, detected with MyoV-CT rabbit antiserum (gift from the Ephrussi Lab). Scale bar: 20µm. (B.) Quantification of intensity line scans around the oocyte cortex for all genotypes overlaid, with data from Figure 6. The localization is similar for wildtype and Capu rescue backgrounds. The posterior cap increase is lost in *capu* null oocytes (yellow line). (B’) Individual genotype scans of cortical localization, average (solid line) with the standard deviation. Wildtype (w[1118]) n= 6, CapuA rescue n = 10, CapuJ rescue n = 10, CapuΔN rescue n = 8, *capu* null = 7. (C.) Quantification of intensity line scans from the anterior to posterior oocyte for all genotypes overlaid. *capu* null data are the yellow line. The anterior/posterior distance was from the nurse cell-oocyte interface to the posterior oocyte cortex-anterior follicle cell border. (C’) Individual genotype scans of anterior/posterior scan, average (solid line) with the standard deviation. Wildtype (w[1118]) n= 6, CapuA rescue n = 10, CapuJ rescue n = 10, CapuΔN rescue n = 8, *capu* null = 7.

**Figure S7.**
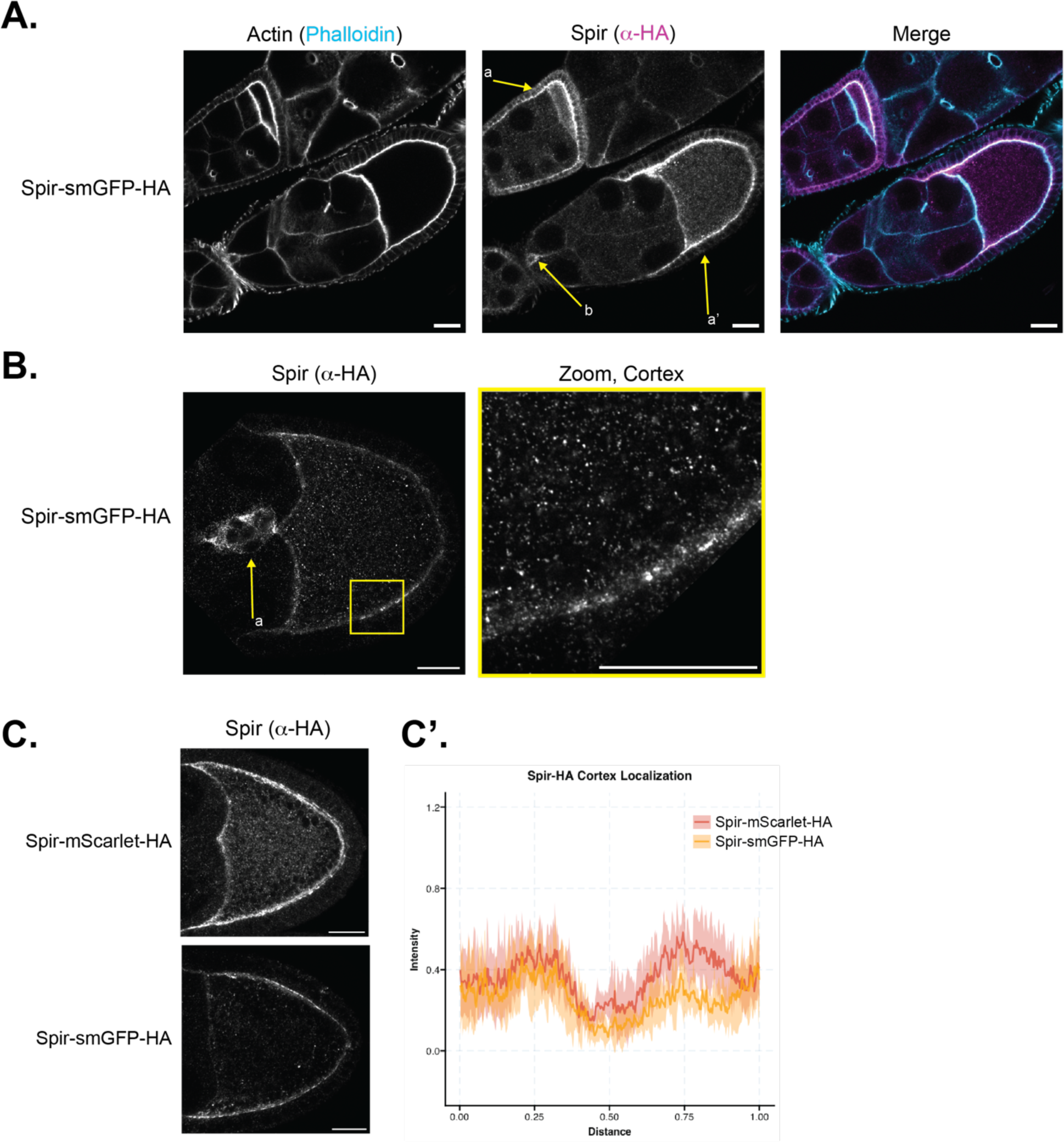
Spir’s localization in the developing egg chamber. (D.) Immunofluorescence of endogenously tagged Spir. Signal is observed in the oocyte, follicle cells (arrows a, a’) and migrating border cell cluster (arrow b). Counterstained with phalloidin to label actin. Scale bars: 20µm. (B.) Immunofluorescent imaging of endogenously tagged Spir in a stage 9 egg chamber using super resolution microscopy. Clear localization is observed in the migrating border cell cluster. Zooming in at the cortex (yellow box) reveals punctate signal along the membrane and in the cytoplasm. Scale bars: 20µm. (C.) Representative images, (Spir-HA, anti-HA 1:1000, CST) and (C’) quantification of Spir signal around the cortex of the oocyte. This shows a dip in intensity at the posterior with both endogenously tagged Spir lines generated. Spir-mScarlet-HA n= 9, Spir-smGFP-HA n = 7.

## Supplementary File 1. smiFISH probes

Lists of probes used to detect *oskar, bicoid, gurken* and FLAP-X complementary sequences.

## Notes

### Competing Interest Statement

The authors have declared no competing interest.

## References

Almonacid, M. and Verlhac, M.-H. (2021). A new mode of mechano-transduction shakes the oocyte nucleus, thereby fine tunes gene expression modulating the developmental potential. C. R. Biol. 343, 223–234.

Alzahofi, N., Welz, T., Robinson, C. L., Page, E. L., Briggs, D. A., Stainthorp, A. K., Reekes, J., Elbe, D. A., Straub, F., Kallemeijn, W. W., et al. (2020). Rab27a co-ordinates actin-dependent transport by controlling organelle-associated motors and track assembly proteins. Nat. Commun. 11, 3495.

Azoury, J., Lee, K. W., Georget, V., Rassinier, P., Leader, B. and Verlhac, M.-H. (2008). Spindle Positioning in Mouse Oocytes Relies on a Dynamic Meshwork of Actin Filaments. Curr. Biol. 18, 1514–1519.

Babu, K., Cai, Y., Bahri, S., Yang, X. and Chia, W. (2003). Roles of Bifocal, Homer, and F-actin in anchoring Oskar to the posterior cortex of *Drosophila* oocytes. Genes Dev. 18, 138–143.

Bence, M., Jankovics, F., Lukácsovich, T. and Erdélyi, M. (2017). Combining the auxin-inducible degradation system with CRISPR /Cas9-based genome editing for the conditional depletion of endogenous *Drosophila melanogaster* proteins. FEBS J. 284, 1056–1069.

Benson, D. A., Cavanaugh, M., Clark, K., Karsch-Mizrachi, I., Lipman, D. J., Ostell, J. and Sayers, E. W. (2012). GenBank. Nucleic Acids Res. 41, D36–D42.

Bor, B., Bois, J. S. and Quinlan, M. E. (2015). Regulation of the formin cappuccino is critical for polarity of *D rosophila* oocytes: Capu is Autoinhibited In Vivo. Cytoskeleton 72, 1–15.

Bose, M., Lampe, M., Mahamid, J. and Ephrussi, A. (2022). Liquid-to-solid phase transition of oskar ribonucleoprotein granules is essential for their function in Drosophila embryonic development. Cell 185, 1308–1324.e23.

Bradley, A. O., Vizcarra, C. L., Bailey, H. M. and Quinlan, M. E. (2019). Spire stimulates nucleation by Cappuccino and binds both ends of actin filaments. Mol. Biol. Cell 31, 273–286.

Calvo, L., Ronshaugen, M. and Pettini, T. (2021). smiFISH and embryo segmentation for single-cell multi-gene RNA quantification in arthropods. *Commun*. Biol. 4, 352.

Cha, B.-J., Serbus, L. R., Koppetsch, B. S. and Theurkauf, W. E. (2002). Kinesin I-dependent cortical exclusion restricts pole plasm to the oocyte posterior. Nat. Cell Biol. 4, 592–598.

Chang, C.-W., Nashchekin, D., Wheatley, L., Irion, U., Dahlgaard, K., Montague, T. G., Hall, J. and St. Johnston, D. (2011). Anterior–Posterior Axis Specification in *Drosophila* Oocytes: Identification of Novel *bicoid* and *oskar* mRNA Localization Factors. Genetics 188, 883–896.

Dahlgaard, K., Raposo, A. A. S. F., Niccoli, T. and St Johnston, D. (2007). Capu and Spire Assemble a Cytoplasmic Actin Mesh that Maintains Microtubule Organization in the Drosophila Oocyte. Dev. Cell 13, 539–553.

Doerflinger, H., Benton, R., Torres, I. L., Zwart, M. F. and St Johnston, D. (2006). Drosophila Anterior-Posterior Polarity Requires Actin-Dependent PAR-1 Recruitment to the Oocyte Posterior. Curr. Biol. 16, 1090–1095.

Doerflinger, H., Zimyanin, V. and St Johnston, D. (2022). The Drosophila anterior-posterior axis is polarized by asymmetric myosin activation. Curr. Biol. 32, 374–385.e4.

Dombrowski, C., Cisneros, L., Chatkaew, S., Goldstein, R. E. and Kessler, J. O. (2004). Self-Concentration and Large-Scale Coherence in Bacterial Dynamics. Phys. Rev. Lett. 93, 098103.

Emmons, S., Phan, H., Calley, J., Wenliang, C., James, B. and Manseau, L. (1995). cappuccino, a Drosophila maternal effect gene required for polarity of the egg and embryo, is related to the vertebrate limb deformity locus. Genes Dev. 9, 2482– 2494.

Ephrussi, A., Dickinson, L. K. and Lehmann, R. (1991). oskar Organizes the Germ Plasm and Directs Localization of the Posterior Determinant nanos. Cell 66, 37– 50.

Ganguly, S., Williams, L. S., Palacios, I. M. and Goldstein, R. E. (2012). Cytoplasmic streaming in Drosophila oocytes varies with kinesin activity and correlates with the microtubule cytoskeleton architecture. Proc. Natl. Acad. Sci. 109, 15109–15114.

Glotzer, J. B., Saffrich, R., Glotzer, M. and Ephrussi, A. (1997). Cytoplasmic flows localize injected oskar RNA in Drosophila oocytes. Curr. Biol. 7, 326–337.

Gutzeit, H. and Koppa, R. (1982). Time-lapse film analysis of cytoplasmic streaming during late oogenesis of Drosophila. J. Embryol. exp. Morph 67, 101–111.

Holubcová, Z., Howard, G. and Schuh, M. (2013). Vesicles modulate an actin network for asymmetric spindle positioning. Nat. Cell Biol. 15, 937–947.

Hudson, A. M. and Cooley, L. (2014). Methods for studying oogenesis. Methods 68, 207–217.

Jaramillo, A. M., Weil, T. T., Goodhouse, J., Gavis, E. R. and Schupbach, T. (2008). The dynamics of fluorescently labeled endogenous *gurken* mRNA in *Drosophila*. J. Cell Sci. 121, 887–894.

Kanca, O., Zirin, J., Garcia-Marques, J., Knight, S. M., Yang-Zhou, D., Amador, G., Chung, H., Zuo, Z., Ma, L., He, Y., et al. (2019). An efficient CRISPR-based strategy to insert small and large fragments of DNA using short homology arms. eLife 8, e51539.

Kerkhoff, E., Simpson, J. C., Leberfinger, C. B., Otto, I. M., Doerks, T., Bork, P., Rapp, U. R., Raabe, T. and Pepperkok, R. (2001). The Spir actin organizers are involved in vesicle transport processes. Curr Biol 11, 1963–8.

Kim-Ha, J., Smith, J. L. and Macdonald, P. M. (1991). oskar mRNA is localized to the posterior pole of the Drosophila oocyte. Cell 66, 23–35.

Krauss, J., López de Quinto, S., Nüsslein-Volhard, C. and Ephrussi, A. (2009). Myosin-V Regulates oskar mRNA Localization in the Drosophila Oocyte. Curr. Biol. 19, 1058–1063.

Leader, D. P., Krause, S. A., Pandit, A., Davies, S. A. and Dow, J. A. T. (2018). FlyAtlas 2: a new version of the Drosophila melanogaster expression atlas with RNA-Seq, miRNA-Seq and sex-specific data. Nucleic Acids Res. 46, D809–D815.

Lee, P.-T., Zirin, J., Kanca, O., Lin, W.-W., Schulze, K. L., Li-Kroeger, D., Tao, R., Devereaux, C., Hu, Y., Chung, V., et al. (2018). A gene-specific T2A-GAL4 library for Drosophila. eLife 7, e35574.

Lu, W., Lakonishok, M., Liu, R., Billington, N., Rich, A., Glotzer, M., Sellers, J. R. and Gelfand, V. I. (2020). Competition between kinesin-1 and myosin-V defines Drosophila posterior determination. eLife 9, e54216.

Lu, W., Lakonishok, M. and Gelfand, V. I. (2023). The dynamic duo of microtubule polymerase Mini spindles/XMAP215 and cytoplasmic dynein is essential for maintaining *Drosophila* oocyte fate. Proc. Natl. Acad. Sci. 120, e2303376120.

Mallart, C., Netter, S., Chalvet, F., Claret, S., Guichet, A., Montagne, J., Pret, A.-M. and Malartre, M. (2024). JAK-STAT-dependent contact between follicle cells and the oocyte controls Drosophila anterior-posterior polarity and germline development. Nat. Commun. 15, 1627.

Manseau, L. J. and Schupbach, T. (1989). cappuccino and spire: two unique maternal-effect loci required for both the anteroposterior and dorsoventral patterns of the Drosophila embryo. Genes Dev. 3, 1437–1452.

Nern, A., Pfeiffer, B. D. and Rubin, G. M. (2015). Optimized tools for multicolor stochastic labeling reveal diverse stereotyped cell arrangements in the fly visual system. Proc. Natl. Acad. Sci. 112, E2967–E2976.

Öztürk-Çolak, A., Marygold, S. J., Antonazzo, G., Attrill, H., Goutte-Gattat, D., Jenkins, V. K., Matthews, B. B., Millburn, G., dos Santos, G., Tabone, C. J., et al. (2024). FlyBase: updates to the Drosophila genes and genomes database. Genetics 227, iyad211.

Parton, R. M., Hamilton, R. S., Ball, G., Yang, L., Cullen, C. F., Lu, W., Ohkura, H. and Davis, I. (2011). A PAR-1–dependent orientation gradient of dynamic microtubules directs posterior cargo transport in the Drosophila oocyte. J. Cell Biol. 194, 121– 135.

Pfender, S., Kuznetsov, V., Pleiser, S., Kerkhoff, E. and Schuh, M. (2011). Spire-Type Actin Nucleators Cooperate with Formin-2 to Drive Asymmetric Oocyte Division. Curr. Biol. 21, 955–960.

Port, F., Chen, H.-M., Lee, T. and Bullock, S. L. (2014). Optimized CRISPR/Cas tools for efficient germline and somatic genome engineering in Drosophila. Proc. Natl. Acad. Sci. U. S. A. 111, E2967–E2976.

Quinlan, M. E. (2013). Direct interaction between two actin nucleators is required in Drosophila oogenesis. Development 140, 4417–4425.

Quinlan, M. E., Heuser, J. E., Kerkhoff, E. and Dyche Mullins, R. (2005). Drosophila Spire is an actin nucleation factor. Nature 433, 382–388.

Ren, X., Housden, B. E., Hu, Y., Roesel, C., Lin, S., Liu, L.-P., Yang, Z., Mao, D., Sun, L., Wu, Q., et al. (2013). Optimized gene editing technology for Drosophila melanogaster using germ line-specific Cas9.

Robinson, D. N. and Cooley, L. (1997). *Drosophila* Kelch Is an Oligomeric Ring Canal Actin Organizer. J. Cell Biol. 138, 799–810.

Rongo, C., Gavis, E. R. and Lehmann, R. (1995). Localization of oskar RNA regulates oskar translation and requires Oskar protein. Development 121, 2737–2746.

Rosales-Nieves, A. E., Johndrow, J. E., Keller, L. C., Magie, C. R., Pinto-Santini, D. M. and Parkhurst, S. M. (2006). Coordination of microtubule and microfilament dynamics by Drosophila Rho1, Spire and Cappuccino. Nat. Cell Biol. 8, 367–376.

Santos, A. C. and Lehmann, R. (2004). Germ Cell Specification and Migration in Drosophila and beyond. Curr. Biol. 14, R578–R589.

Scheffler, K., Uraji, J., Jentoft, I., Cavazza, T., Mönnich, E., Mogessie, B. and Schuh, M. (2021). Two mechanisms drive pronuclear migration in mouse zygotes. Nat. Commun. 12, 841.

Schuh, M. and Ellenberg, J. (2008). A New Model for Asymmetric Spindle Positioning in Mouse Oocytes. Curr. Biol. 18, 1986–1992.

Serbus, L. R. (2005). Dynein and the actin cytoskeleton control kinesin-driven cytoplasmic streaming in Drosophila oocytes. Development 132, 3743–3752.

St. Johnston, D., Beuchle, D. and Nusslein-Volhard, C. (1991). staufen, a Gene Required to Localize Maternal RNAs in the Drosophila Egg. Cell 66, 51–63.

Stürner, T., Ferreira Castro, A., Philipps, M., Cuntz, H. and Tavosanis, G. (2022). The branching code: A model of actin-driven dendrite arborization. Cell Rep. 39, 110746.

Tanaka, T., Kato, Y., Matsuda, K., Hanyu-Nakamura, K. and Nakamura, A. (2011). Drosophila Mon2 couples Oskar-induced endocytosis with actin remodeling for cortical anchorage of the germ plasm. Development 138, 2523–2532.

Thielicke, W. and Sonntag, R. (2021). Particle Image Velocimetry for MATLAB: Accuracy and enhanced algorithms in PIVlab. J. Open Res. Softw. 9, 12.

Thielicke, W. and Stamhuis, E. J. (2014). PIVlab – Towards User-friendly, Affordable and Accurate Digital Particle Image Velocimetry in MATLAB. J. Open Res. Softw. 2,.

Tittel, J., Welz, T., Czogalla, A., Dietrich, S., Samol-Wolf, A., Schulte, M., Schwille, P., Weidemann, T. and Kerkhoff, E. (2015). Membrane Targeting of the Spir·Formin Actin Nucleator Complex Requires a Sequential Handshake of Polar Interactions. J. Biol. Chem. 290, 6428–6444.

Trovisco, V., Belaya, K., Nashchekin, D., Irion, U., Sirinakis, G., Butler, R., Lee, J. J., Gavis, E. R. and St Johnston, D. (2016). bicoid mRNA localises to the Drosophila oocyte anterior by random Dynein-mediated transport and anchoring. eLife 5, e17537.

Tsanov, N., Samacoits, A., Chouaib, R., Traboulsi, A.-M., Gostan, T., Weber, C., Zimmer, C., Zibara, K., Walter, T., Peter, M., et al. (2016). smiFISH and FISH-quant – a flexible single RNA detection approach with super-resolution capability. Nucleic Acids Res. 44, e165–e165.

Vanzo, N. F. and Ephrussi, A. (2002). Oskar anchoring restricts pole plasm formation to the posterior of the *Drosophila* oocyte. Development 129, 3705–3714.

Venken, K. J. T., Schulze, K. L., Haelterman, N. A., Pan, H., He, Y., Evans-Holm, M., Carlson, J. W., Levis, R. W., Spradling, A. C., Hoskins, R. A., et al. (2011). MiMIC: a highly versatile transposon insertion resource for engineering Drosophila melanogaster genes. Nat. Methods 8, 737–743.

Vizcarra, C. L., Kreutz, B., Rodal, A. A., Toms, A. V., Lu, J., Zheng, W., Quinlan, M. E. and Eck, M. J. (2011). Structure and function of the interacting domains of Spire and Fmn-family formins. Proc. Natl. Acad. Sci. 108, 11884–11889.

Weil, T. T., Forrest, K. M. and Gavis, E. R. (2006). Localization of bicoid mRNA in Late Oocytes Is Maintained by Continual Active Transport. Dev. Cell 11, 251–262.

Zimyanin, V. L., Belaya, K., Pecreaux, J., Gilchrist, M. J., Clark, A., Davis, I. and St Johnston, D. (2008). In Vivo Imaging of oskar mRNA Transport Reveals the Mechanism of Posterior Localization. Cell 134, 843–853.

